# A topological method of generating action potentials and EEG oscillations in a surface network

**DOI:** 10.1101/2022.06.25.497598

**Authors:** Siddhartha Sen

## Abstract

The brain is a source of continuous electrical activity, which include one dimensional voltage pulses (action potentials) that propagate along nerve fibres, transient localised oscillations, and persistent surface oscillations in five distinct frequency bands. However, a unified theoretical framework for modelling these excitations is lacking. In this paper we provide such a framework by constructing a special surface network in which all observed brain-like signals, including surface oscillations, can be generated by topological means. Analytic expressions for all these excitations are found and the values of the five frequency bands of surface oscillations are correctly predicted. It is shown how input signals of the system produce their own communication code to encode the information they carry and how the response output propagating signals produced carry this input information with them and can transfer it to the pathways they traverse as a non-transient topological memory structure of aligned spin-half protons. It is conjectured that the memory structure is located in the insulating sheaths of nerve fibres and are stable only if the pathways between assembly of neurons, that represents a memory structure, includes loops. The creation time and size of memory structures is estimated and a memory specific excitation frequencies for a memory structure is identified and determined, which can be used to recall memories.

## 1 Introduction

In this paper we introduce a special topological surface network with surface electrons and protons that captures a key global feature of the brain. An analysis of the network shows that it can generate all brain-like signals, store memories as aligned proton spin structures located in the insulating myelin sheaths of nerve fibres.

The brain consists of a vast number (*N* ≈ 10^11^) of neurons [1]. Each neuron has multiple signal receiving stations, protrusions called dendrites. Signals entering the dendrites are processed as they enter the neuron, and if they cross a certain threshold energy, a processed signal or action potential, is generated that exits the neuron, as a one dimensional voltage pulse via its one single signal outlet nerve fibre, the axon. The axon terminal of one neuron passes on the signal it carries to the dendrite of another neuron in the form of a cloud of specialized molecules called neurotransmitters. There is a gap of about 10^−6^*cm* between the axon of one neuron and the receiving dendrite terminal, of another neuron, called a synapse. The membranes that cover the neuron and axons contains ions and a voltage difference between the inner and outer surface of the membranes is observed. Protein gates on the axon membrane connect its inner and outer surface and open and close under appropriate conditions to allow ions to flow between them. It is believed that electrical signals generated by the brain are due to ion flows using properties of these ion gates[1].

A variety of electrical activities are observed in the brain. They include different types of one dimensional voltage pulses that propagate along the axons, localized transient oscillations, as well as the ever present brain surface voltage oscillations revealed by Electroencephalography (EEG) measurements[2] on the scalp. The EEG oscillations are observed to belong to five frequency bands, called, the delta (.5-3 Hz), theta (3-8 Hz), alpha (8-12 Hz), beta (12-30 Hz), and gamma (30-42+ Hz) waveforms. It is established that action potentials play an essential role in the functioning of the brain, they initiate motor or emotional responses, while EEG waveforms seems to respond whenever action potentials are generated but the precise link between them is not clear.

Understanding the autonomous functioning of this vast and complex system is a formidable task. But remarkable progress in unravelling key features of the way the brain functions have been made. Observational neuroscience is progressing at a rapid pace using a variety of innovative experimental techniques to suggest how memories are created, how they may be manipulated, how they may be stored in collections of special engram cells[3, 4, 5], and how memories may be space and time tagged[6, 7]. Theoretical neuroscience too is progressing [8] driven by new ideas of theoretical representations of the brain[9] and increased computational power. Methods to theoretically model behaviour, to discover new excitations, suggested ways to understand complex brain events and store memory[10] are emerging[11].

However there are major conceptual theoretical problems that remain unaddressed. Areas of concern include the lack of theories that allow signals to carry information[1], the lack of a theoretical understanding of how memories are stored[12] and the lack of understanding the properties of observed EEG waveforms are generated[2].

Observations strongly suggest that sensory input signals to the brain carry with them information that is processed to give us our sensory experiences. If the brain regards that some incoming information is important then it is stored as long term memory. The process of memory creation has been very well studied[13] but where they are stored is still under investigation. A current suggestion is that memories may be stored in the pathways between special engram cells[4]

Interactions between incoming signals and memory are essential for our brain to function. Such interactions allow us to identify people we know, to avoid dangerous places, help us to remember and return to where we live and to engage in conversations. Yet theoretical neuroscience cannot address this issue. In current theoretical neuroscience memory information is fed into the system by special input signals and retrieved by special input cues. They are not directly related to all incoming input signals. Brain functions are currently modelled by introducing interaction between individual or populations of neurons that are either always excitatory or inhibitory in linear networks that are chosen to reflect the observed connectivity of the brain or that describe important pathways between brain organs. Simulation of brain activities are carried out by introducing by “integrate and fire” input signals[8] that do not and cannot carry biological information. Thus two important features of the brain are ignored. Signals do not carry information so that no theoretical link between signals and memory is possible and the constant feedback interactions and information exchange between signals and memories that occurs in the brain are missing.

Another major theoretical gap is the absence of understanding the origin of EEG waveforms and the way they interact with other brain signals. A current suggestion[2] that EEG waveforms are created by dipole current loops on dendrites cannot explain the observed properties EEG waveforms. This gap means that there is no theoretical way to describe the interaction of EEG waveforms with memory that are observationally seen to occur[33].

We will prove that current theoretical signals cannot carry information. This means that even if the integrate and fire signals are replaced by realistic signals, such as the membrane ion flow model of Hodgkin Huxley[1] signals or the acoustic membrane signals of Jackson and Heimburg[14] they still would not provide the information loop between signals and memory required to understand cognitive functions of the brain as even these signals cannot carry information.

Let us prove this result for the Hodgkin Huxley[1] ion membrane model and the membrane acoustic soliton model[14]. The ion membrane model, as pointed by Scott[1], is a dissipative wave model. In it inflows and outflows of ions between the inner and outer membrane surfaces of an axon leads to the injection of energy in the centre of the axon tube in the form of a voltage gradient. The voltage gradient produced is an energy source and produces a dissipative ohmic current. The process of ion flows is triggered when an incoming signal crosses a voltage threshold that leads to the opening of membrane gates. No other condition is necessary. Thus the signals produced propagate by following a causal cycle of energy injection and dissipation. Such a dissipative wave cannot carry information regarding their creation as causal cycles of energy injection and dissipation that depend only on membrane properties and not on the initial cause of the signal. In modelling the process of energy injection the membrane’s inner and outer surface ion distributions are found by using thermodynamics. But thermodynamic results are independent of the history of how the system parameter values are reached. The conclusion is that signals in dissipative wave theories follow a causal cycle that does not allow signals produced to carry information about their creation. The observed link between signal information and memory is missing in such theories. Scott, who was a distinguished applied mathematician, compares nerve signal propagation with the dissipative mechanism of a burning candle as discussed by the great scientist Faraday in his Christmas lectures [1]. The analogy is that a burning candle is also a dissipate process in which melted wax reaches the tip of the wick by capillary action, becomes a vapour and injects energy that is dissipated by the burning flame. This causal cyclic dissipative process of the burning candle carries no information regarding how the process was started. It is a coherent process that continues as long as the energy source and the dissipative processes are operative. It has no memory of how the process was started.

The acoustic soliton approach of Jackson[14]unfortunately faces the same problem. Acoustic solitons discovered are non dissipative, non topological excitations that are produced as membrane acoustic waves. They depend on the axon membrane’s elastic properties and the fact that the fluid inside the axon undergoes a phase transition when compressed. Thus a key step for producing soliton signals involves thermodynamic properties and are thus independent of their history so the soliton waves produced cannot carry information of their creation.

These results impact on the capabilities of theoretical approaches to understand memory formation and place limitations on current approaches to model cognitive brain functions [15, 16, 17].

Thus a radical deviation from existing approaches is needed. Our aim is to show that there is such an approach. This is demonstrated by constructing a theoretical surface network which creates its own communication code, produces the range of brain excitations observed, including EEG waveforms, where signals generated clearly carry information regarding their creation, and a mechanism, based on the laws of physics, is suggested for the signals to transfer the information they carry to form stable memories. In the network memories are retrieved by a resonance excitation method, as it is shown that the memory structure created has a memory specific excitation frequency, signals produced can interact with each other and they have information exchange loop with memory. But the method of signal generation suggested is unconventional.

### 1.1 Topological Ideas and Riemann Surfaces

The stated properties of the special surface network properties come from its topological features. Topology is a mathematical discipline in which objects that can be deformed into each other by smooth deformations are regarded as the same. Thus a soccer ball and a rugby ball are topologically identical.^*^.

The surface network we chose has topological properties. It exactly capture the topological connectivity of the axons of the brain as a surface. We will prove this result. It can produce all brain-like signals in response to local topology changing deformations of a subunit, only if the surface is described by a specific algebraic equation that uniquely defines a special Riemann surface. Riemann surfaces provide a geometric description of any polynomial equation in two complex variables.[19]. Thus a special polynomial equation is represented by a special Riemann surface. We will explain why a special Riemann surface is required for signal generation. The final property of the surface is that it must have charged surface particles of spin-half.

The presence of surface spin half particles introduces an additional essential layer of topology to the surface. Without surface spin-half particles none of our results would hold. The topology due to spin-half particles comes from the ability of such particles to arrange themselves in topological distinct patterns because spin-half particles have magnetic properties.

The topological structure due to spin is called a spin structures.[24] The topological connectivity of the surface network can be described by a integer *g* called the genus of the Riemann surface while its spin topological structure can be described by another integer *W* called its topological spin number. These numbers will be defined shortly. Signals are generation by local topology changing surface deformations of subunits of the Riemann surface that deform the topology numbers (*k, w*), that describe the subunits connectivity and spin structure, to the values (*k* = 0, *w* = 0) which describes a spherical surface. Such a process generates *k* non-dissipative one dimensional voltage pulses, called solitons.

### 1.2 A surface connectome is a Riemann surface

We now prove that an exact representation of the topological connectivity features of the brain’s axons by a mathematical Riemann surface is possible. Consider a hypothetical brain connectome^†^, that describes the axon connectivity features of a brain as a one dimensional network, embedded in three dimensions, where each neuron is a nodal point of the network and the lines are axons. The network is enormously complicated and unknown for the human brain. We now replace each line of the connectome by a tube and each nodal point by a suitable nodal surface Fig(1).

**Figure 1:**
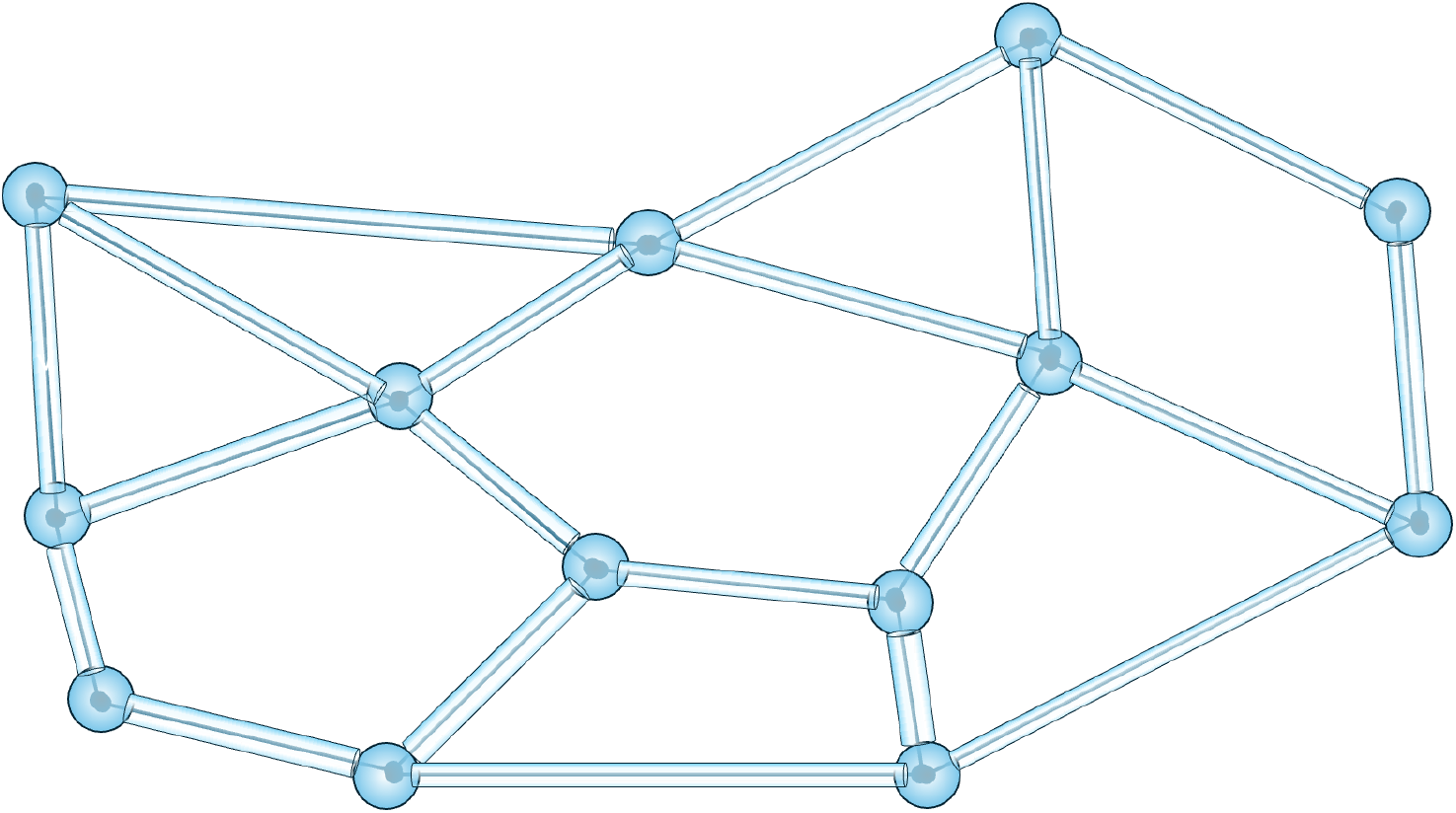
A part of a brain connectome represented by a surface, where the lines of the connectome are replaced by tubes and junction points (neurons) are replaced by junction spheres. See text for details.

This step seems to make the original network even more complex but a remarkable result from the mathematical discipline of topology[31] now comes to our aid. The topological result tells us that any closed surface, no matter how intricate or complex, is topologically equivalent to a sphere with *g* handles attached to it (Fig 2) or an orderly collection of *g* doughnuts[32] (Fig3).

**Figure 2:**
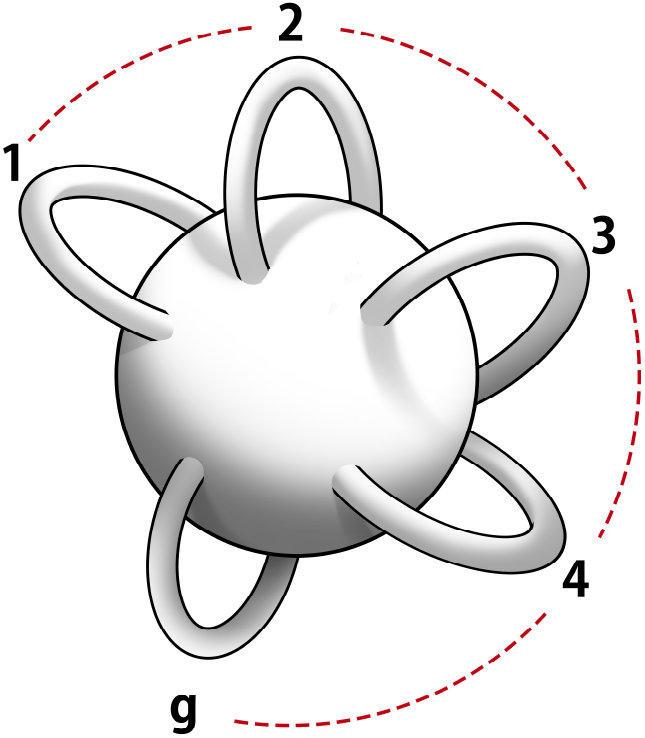
Genus *g* surface. For g=5

The only topological difference between any two surfaces is in the value of one integer number *g* called its genus. This number, the genus *g*, completely captures the topological connectivity of a surface. If we assume, as we do, that the surface is smooth then it can be represented by a mathematical surface known as a Riemann surface of genus *g*.

Thus from the classification theorem it follows that the topological connectivity of any brain connectome can be exactly represented as a smooth Riemann surface of unknown genus *g*, and thus is a geometric representation of an algebraic equation.

### 1.3 Signal Generation and the Riemann theta function

We next explains how brain signals can be generated by topological means and then sketch why such signals are solutions of the non-linear Schroedinger equation. A topological way to produce soliton pulses was discovered by [20] Mumford, who showed that soliton solutions of the non linear KdV differential equation could be generated by local topology changing deformation of a Riemann surface with spin structure.

We need to modify Mumford’s ideas in two ways so that the approach can produce the range of brain-like we are interested in and at the same time coproduce EEG like waveforms. These requirements can be met by an extension of Mumford’s idea, encapsulated as the dynamical law of the network.

The second modification introduced is a restriction. Thus, unlike Mumford, the network surface, cannot be an arbitrary Riemann surface, but must be one that represents a specific algebraic equation known as hyperelliptic equation which is defined by the equation 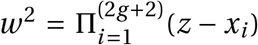 where the roots of the polynomial are either real or appear as a complex conjugate pairs. Recall if *z* = *a* + *ib*, the 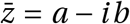 are a complex conjugate pair. The surface must also have charged spin-half particles, such as electrons and protons. This restriction allows the system to produce the wide variety of brain-like signals we want. The mathematical reason for this restriction is explained in Arberello[21].

The surface network can be visualised as a membrane cover of the individual neurons of the brain with the topological connectivity of the brain’s connectome. Thus the Riemann surface tubes are axons, its nodal junction regions are the location of neurons and the points where the tubes enter a junction are the location of dendrites. We will use these descriptive terms.

### 1.4 Input Pinch deformations and EEG waveforms

The first step is to explain how the topology of a subunit is changed by local input signals, called pinch deformations. Pinch deformations reduce the circumference of an axon tube at a point to zero and thus change the topology of the system. Consider a subunit of genus *k* < *g* and spin structure *w* < *W*. Pinch deformations of this subunit reduce its topology numbers from (*k, w*) to (0, 0). The topology numbers (*k* = 0, *w* = 0) as stated before, describe a topological sphere. When this happens we will see, *k* one dimensional soliton voltage pulses that exit through the the axons that connect the subunit to other subunits of the surface and each soliton pulse carries with it a topological spin number phase *e*^*iπw*^. The excitations carry away the topology numbers of the subunit and as well as the pinch deformation that produced them. Thus signals carry information encoded in the system’s own communication code given by pinch deformation parameter values. We have yet to define the spin-topology number *w*. This will be done shortly.

The process described implies that whenever signals are produced by pinch deformations topological spheres are created. It is expected these topological sphere surfaces will have voltage oscillations. We identify these spheres as an individual neuron surfaces.^‡^ and the surface oscillations of an assembly of neurons as the source of the observed scalp EEG waveform voltage oscillations.

### 1.5 Signals are Solutions of a non-linear equation

We next explain the surprising result that all signals generated by the pinch deformation of subunits must be solutions of a special non-linear differential equation called the non-linear Schroedinger equation.

To do this we first introduce an appropriate function to describe the network’s response to pinch deformations. Recall if we are interested in the oscillations of a circle we introduce sine and cosine functions that have as variables the angle of the circle. For a Riemann surface there is an analogous function called the Riemann theta function. The variables that define it depend on, as expected, on the variables of the Riemann surface. Unlike the sine and cosine functions that depend on the one variables that defines a circle, the number of variables in the Riemann theta function is more than what is necessary. This means the variables cannot be freely chosen. The Riemann theta function has to satisfy constraints. A mathematical result tells us that the constraint is a non linear identity.

We now have the concepts required to state the dynamical law for the network. It requires that both input pinch deformations and the responses they produced, described in terms of Riemann theta functions must be compatible with the Riemann surface structure even in the pinch limit. It is then shown that the non-linear constraints, in the limit of a pinch deformation, become a set of non linear Schroedinger equations, only if it represents a special algebraic equation, then the response to pinch deformations are Riemann theta functions that satisfy the non-linear Schroedinger differential equation. They exist and their analytic forms are known. The technical details are omitted[20]. We now postulate the dynamical law of the network. The dynamical law has two parts. The first part notes that all deformations of the Riemann surface are automatically transferred to the deformation of the variables of the Riemann theta function associated with it as these are related, explained already.

The second part makes use of a mathematical theorem that states that a Riemann theta function represent a Riemann surface only if it satisfies a non linear identity called the Fay trisecant identity[22, 20]. We will explained the intuitive basis of this requirement. A more technical argument will be sketched later.

The dynamical law thus requires that when the Riemann surface is deformed, the Riemann theta function with their variables automatically deformed, must continue to satisfy the Fay identity. Since in the pinch deformation limit the Fay identities[23]become a set of non linear Schroedinger equations[23] the Riemann theta responses to pinch deformations must be solutions of the nonlinear Schroedinger equation. But the non-linear Schroedinger result holds only when the Riemann surface represents a hyperelliptic equation.

Thus the dynamical law tells us that input pinch deformations produce outputs that are solutions of the non linear Schroedinger equation. Solutions of the non linear Schroedinger equation can represent a wide variety, possibly all, observed brain like signals and their analytic forms are known and the solutions carry pinch deformation information. Thus the claim is that any given observed brain like signals, no matter how it is produced, can be fitted by choosing suitable pinch deformation parameters. We will give an example of such a fit in a later section.

### 1.6 How Signals Produce Memory

We sketched why have voltage soliton pulse signals produced by pinch deformations are described in terms of the Riemann theta function, where the Riemann theta function variables contain in them pinch deformation information and they are solutions of the non-linear Schroedinger equation.

We now show how these transient soliton signal voltage pulse signals can transfer the information they carry to form non transient memory structures of aligned surface spin-half particle, in the pathways they traverse. This information transference process happens because of the laws of electromagnetism. Moving pulses carry charge and thus generate a helical transient magnetic field[28],^§^which acts on the magnetic spin half surface particles, aligning them to form a non transient structure that captures the magnetic field profile and hence contains in it all deformation details responsible for created the voltage pulse signal.

This helical spin structure created, if stable, is a substrate for memory.^¶^ We show the structure has a signal specific excitation frequency. This means that long term memories can be recalled by a resonance excitation method by a suitable excitation frequency. We will, in a later section, discuss the requirements to create stable memories and derive a formula for their creation time and size. We have given an intuitive account of how the surface network can produce signals and store memory. Our next step is to add in the missing mathematical details.

## 2 Outline of Paper

First we give an outline of the paper. After that we address mathematical issues. In the next section the mathematical variables that define a Riemann surfaces and its associated Riemann theta functions are introduced and the spin topology number *W* is defined. We then explain why the Riemann theta function has to satisfy the Fay trisecant identity at a more technical level. This is followed by postulating a dynamical law of the network in formal terms. We then define the input signals of the network as local pinch deformations. ^||^ The analytic form of moving voltage soliton pulse signals are known. This means we can determine the helical magnetic fields they generate and then from standard methods of physics, determine the spin aligned memory structure of spin-half surface particles in response to the transient magnetic field of the moving solitons. We are able to theoretically show that each memory structure will have its own characteristic excitation frequency its excitation frequency and determine the frequency values they have. We then estimate the the time required to create memory structures. The creation time estimate comes from the conditions needed for the memory structure to be stable. We then discuss how memories can be retrieved by exploiting their excitation frequencies.

After that we then turn to discuss EEG waveforms generation. We show that they can be identified as sphere surface voltage oscillations that are produced whenever pinch deformation generated signals are generated by pinch deformations. We theoretically predict that these oscillations have five oscillation frequency bands with amplitude values inversely related to their frequencies.

Following these general results we shift our attention to a special subclass of tiling solutions of the wave equation known as dihedral tiling. These solutions are special as they form a complete set of solutions. There are an infinite number of them and linear combinations of these solutions can be used to represent all other waveforms as well as arbitrary function on the surface of the sphere. For the rest of the paper we restrict ourselves to these solutions and use their special mathematical features to develop the mathematical tools required to analyse the interaction of EEG waveforms with other brain excitations and with memory. With these theoretical results in place we proceed to model and explain a sequence of brain excitations that are observed during deep sleep.

We end by listing the testable predictions of the approach, drawing attention to some of its special features that might provide a different way to understand brain functions, and outlining future work. Our most significant prediction is that memories are encoded in aligned helical surface spin-half protons located in the insulating myelin sheaths of axons. We provide evidence that support the conjecture. We also predict that a topologically stable memory structure is an assembly of neurons connected together by axon pathways that have non-zero spin topology number. This means the axon pathways must have loops. The predicted memory structure is similar to an engram.

Let us define the variables we need.

## 3 Riemann Surfaces

### 3.1 Riemann Surface and theta function variables

A Riemann surface is a smooth topological surface. It is defined by topological coordinates that describe its connectivity and a mathematical object, called a one form, that describes its smoothness. We define these coordinates and the smoothness measure. We then show how these Riemann surface variables are used to define the variables of its associated Riemann theta function.

As a Riemann surface represents an algebraic equation hence it is expected that it’s topological coordinates and it’s smoothness measure must be constructible from its associated algebraic equation, in our case the hyperelliptic equation.

We will briefly indicate how this can be done. This sketch makes the conceptual features of the approach clear.

The topological connectivity of the Riemann surface is captured by 2*g* loops, as shown in Fig 3. They are *g* loops *a*_*i*_ that go round the tubes of the Riemann surface as shown while the *g* loops *b*_*i*_ circle round loops of the doughnuts, as shown. The smoothness properties of a Riemann surface are captured by a set objects known as a one form. A one form can be written locally as *f* (*z*)*dz*, where *z* is a complex number that represents a surface point. It is an an object that can be integrated along a line or loop. Riemann proved that a genus *g* Riemann surface has exactly *g* smooth one forms. We write them as *ω*(*z*)_*j*_ *dz* where the functions *ω*(*z*)_*j*_ are smooth functions, where *z* is a complex variable that represents a point on the genus *g* Riemann surface. These one forms can be integrated over the loops (*a*_*i*_, *b*_*i*_) of Fig 3, to give complex numbers. Riemann normalised these one forms so that 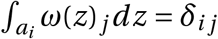 where *δ*_*ij*_ = 1, when *i* = *j* but is zero otherwise. Riemann then proved that 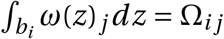 was a complex symmetric matrix and that its imaginary part entries are all positive. It is called the period matrix of the Riemann surface. This ends our mathematical description of a Riemann surface.

**Figure 3:**
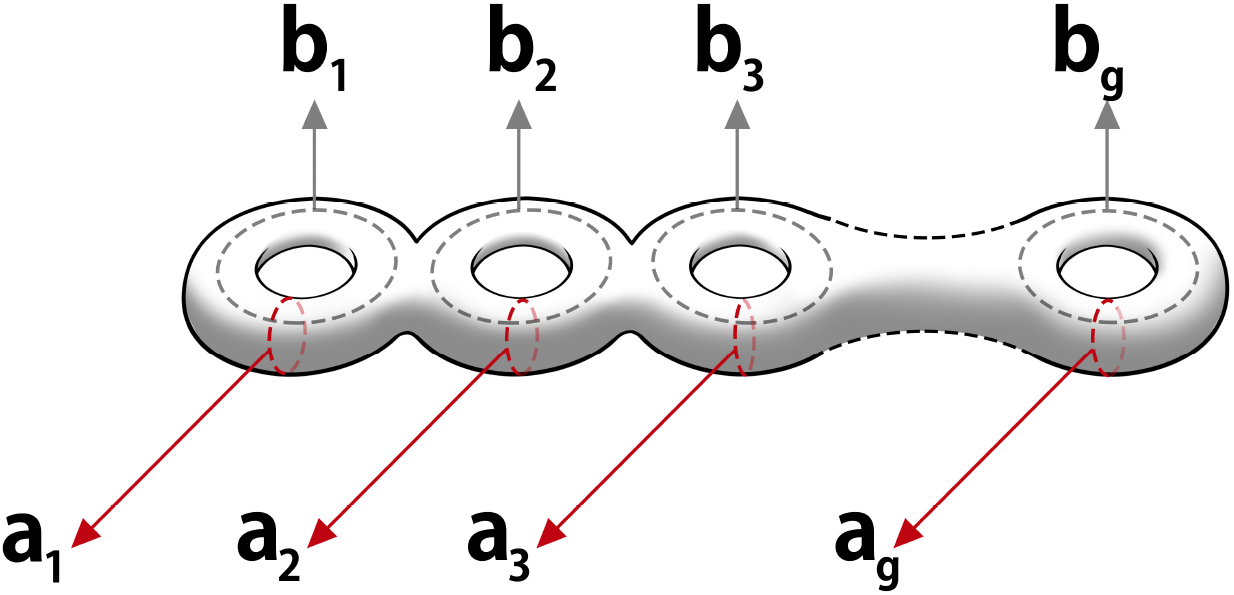
Marked Riemann Surface with loop coordinates

In our discussions we consider a special Riemann surface of genus *g* that represents the hyperelliptic equation 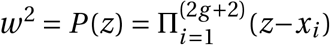. We now briefly sketch how to construct the coordinates and one forms of this genus *g* Riemann surface starting from the equation. The *g* one forms are defined by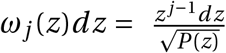 and the 2*g* loops can be constructed from the location of the (2*g* + 2) zeros of *P* (*z*). The details of the construction are in Mumford.

We next define the algebraic Riemann theta functions. Its variables are constructed from the loops and one forms of its associated Riemann surface and it also includes the spin structure of its Riemann surface. The spin structure variables in the Riemann theta function are defined by a set of 2*g* discrete parameters called the characteristics of the Riemann theta function. These parameters (*α*_*i*_ .*β*_*i*_) are associated with the (*a*_*i*_, *b*_*i*_) loops of the Riemann surface and each one of them is either zero or half. It was proved by Mumford[20] that they capture the spin structure of its associated Riemann surface. We now define the Riemann theta function with characteristics that is associated with a genus *g* Riemann surface, 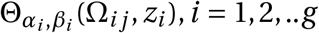 iby the expression

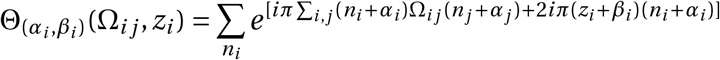

The variables (*z*_*i*_, Ω_*ij*_) are constructed from the Riemann surface variables as explained, while the variables (*α*_*i*_, *β*_*i*_) capture, as stated, the spin structure of the Riemann surface. They are discrete variables are called characteristics. Each of them can be either zero or half, each integer *n*_*i*_ variable ranges between (±∞). The characteristics obey the following algebraic rules: *nα*_*i*_ = *nβ*_=_0, if *n* is an even integer and *nα*_*i*_ = *α*_*i*_, *nβ*_*i*_ = *β*_*i*_ when *n* is an odd integer. where multiplication by *n* means going round the *a*_*i*_ loop *n* times for *α*_*i*_ and for going round the *b*_*i*_ loop *n* times for *β*_*i*_.

The additional set of *g* complex numbers *z*_*i*_ are defined by 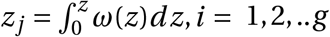 where *z* is an arbitrary point of the Riemann surface. Thus a point on the Riemann surface produces *g* points in its Riemann theta function. We can now define our spin topology number 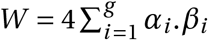. It is an integer and will play an important role in fixing the nature of EEG waveforms to be tiling oscillations.

We return to define a pinch deformation. Geometrically, it is a local topology changing deformation in which the *a*_*i*_ loop circumference shrinks to zero radius at a point,(Fig 4). We capture this geometric picture in an algebraic way as follows. The integral of a one form *ω*_*j*_(*z*)*dz* on the loop *a*_*i*_ can be evaluated the integral is 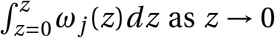 as *z* → 0. The evaluation can be carried out by expanding *ω*_*j*_(*z*)*dz* near *z* ≈ 0 as a Taylor expansion of *ω*_*j*_(*z*)*dz*. The expansion coefficients define the distortion parameters. They will depend on the mathematical properties of the Riemann surface. A careful discussion of how this is done is explained in a separate work[2].

**Figure 4:**
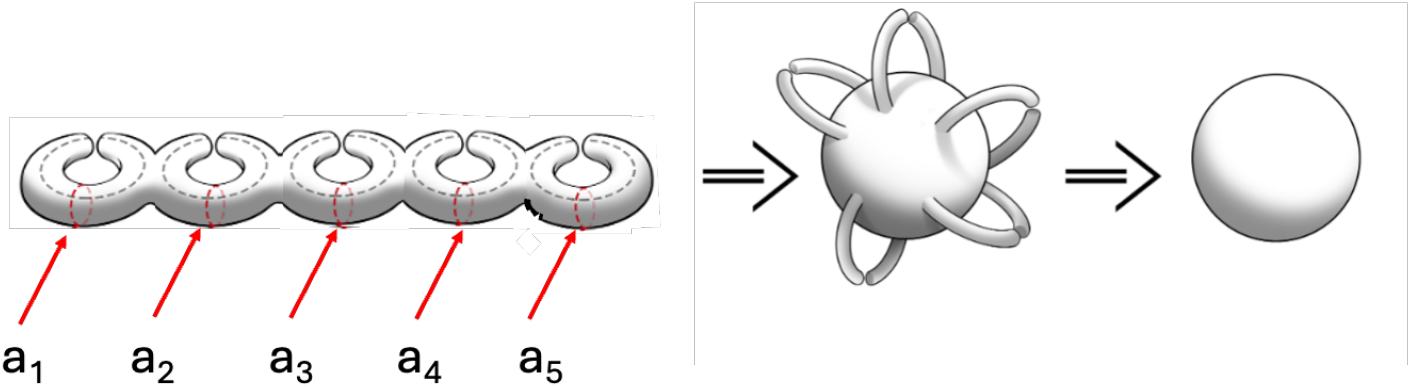
Degenerate Riemann Surface due to pinching. The first figure has *j* doughnuts with *j* = 5 tubes pinched, the second shows the same pinching now represented as 5 handles pinched, the final figure shows that these pinched systems are topological spheres.

We next explain why the Riemann theta function has to satisfy a constraint for it to represents a Riemann surface.

### 3.2 The Fay identity and the Dynamical Law

It is known that the nature of a given Riemann surface of genus *g* is fixed by (6*g* − 6) real parameters called its moduli [20]. These parameters capture the shape of the Riemann surface. However, the period matrix of a Riemann theta function has *g* (*g* + 1) real parameters.[20]. For *g* > 2 the Riemann theta period matrix thus has more parameters than is required. This means that an arbitrary period matrix need not represent a Riemann surface. The parameters of the period matrix are not independent but have to satisfy constraints. But a mathematical result proves that if the Riemann theta function satisfies a constraints called the Fay trisecant identity[20] then the theta function will represent a Riemann surface. The Fay identity only holds if the Riemann surface has surface spin half particles[22]. Hence the presence of particle with spin half, on our surface network, turns the constraint problem to a well defined requirement. We can now state the dynamical law,

#### Both input distortion signals and the surface responses they generate must preserve the Riemann structure of the surface

Thus the dynamical law requires the Riemann theta function must satisfy the Fay trisecant identity as this ensures that it continues to represent its associated Riemann surface even during deformation. In the pinch limit the non linear Fay constraint becomes a set of non linear Schroedinger equations[23], provided the Riemann surface represents a hyperelliptic equation. For Riemann surfaces defined by other algebraic equations other non-linear differential equations emerge. Hence we have the surprising result that for our Riemann surface, local pinch deformation generate excitations that are solutions of the non-linear Schroedinger differential equation. This result follows from the dynamical law. The technical details are not important for our discussions and are omitted. They are given in Kalla[23]. Later we will use the dynamical law to calculate properties of EEG waveforms. There the requirement of preserving the underlying Riemann surface structure is enforced by insisting that only results that are conformal invariant, which means that have local scale invariance, are acceptable, since these transformations preserve Riemann surface structure[19]. We have stated that soliton solutions and other excitations, created by pinch deformations, can produce brain-like signals. We now provide some observational evidence in support of this claim. Two sets of spike trains of action potential pulses were produced from a thalamic neuron of mice, one with two and the other with five spikes. These results were then fitted to the N-soliton solution of the non-linear Schroedinger equation found by Previato[27]. The fits first select the number *N* of the solution, and then for the chosen *N* solution, the deformation parameter values, present in the solutions, are adjusted to fit the observational data. We reproduce these fits from our paper[2], where the data and fit details are given. The fits are very good. This result supports the claim that action potentials can be represented by solitons. Previato’s solution is written down in the Supplementary section.

Let us present the examples. The same thalamic neuron in a mouse, as stated, is excited in two different ways. Once by presynaptic simulation, and then by the injection of square voltage spikes. The data curves are in blueand the fits are in red (Fig5 and Fig6).

**Figure 5:**
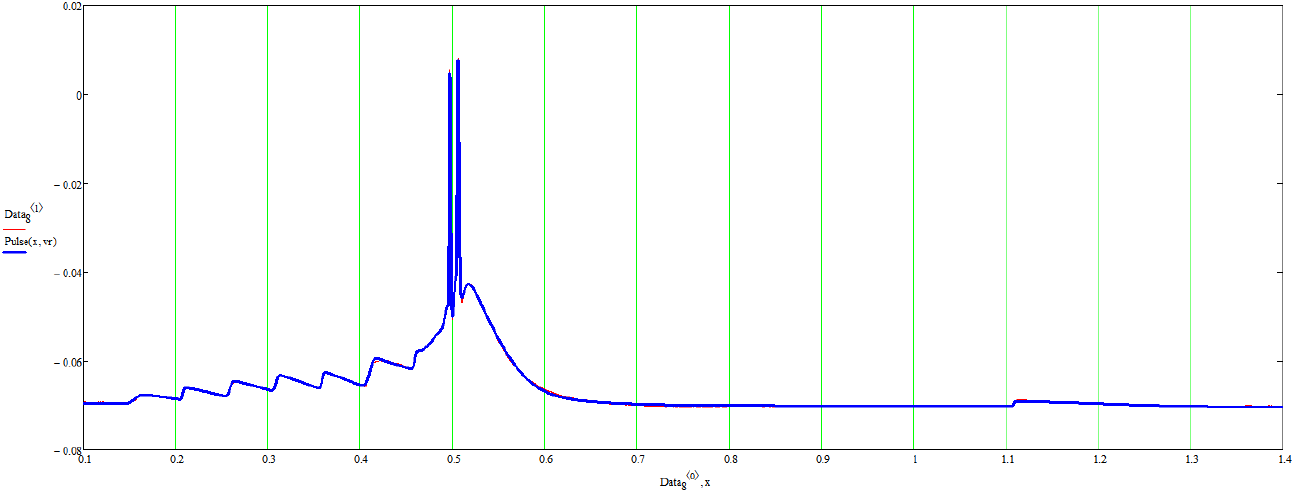
A firing thalamic neuron triggered by presynaptic simulation(blue) and the fit (red). See text for details.

**Figure 6:**
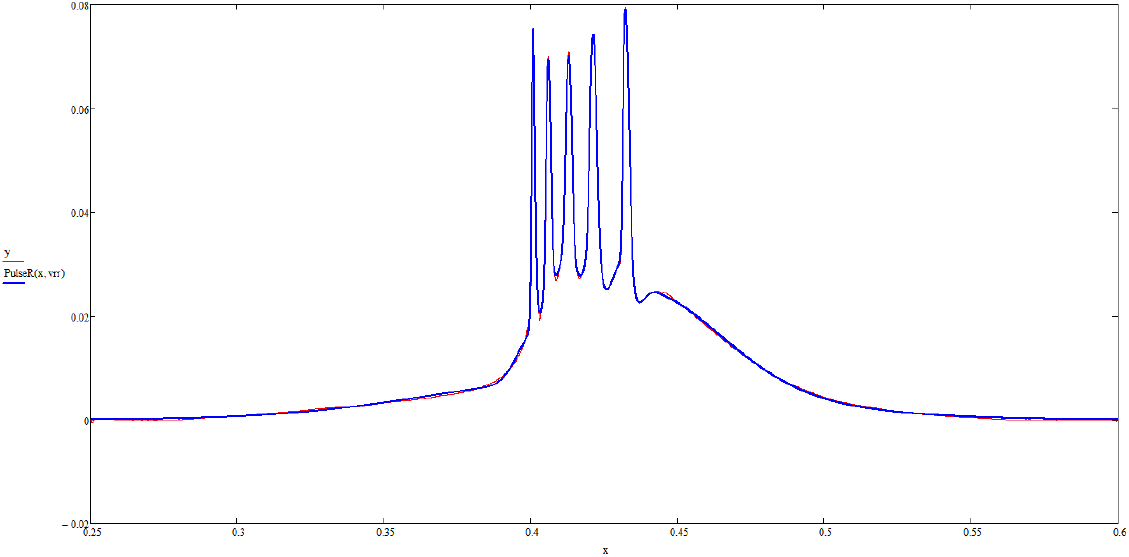
A firing thalamic neuron triggered by the injection of square voltage pulses(blue) and the fit(red). See text for details.

The example shows that multi-soliton solutions can indeed fit observed data and that signals produced in different ways can still be fitted by suitable pinch deformation. Finally each pulse of the spike train, is found to be different, as theoretically predicted.

Let us next turn to discuss features of soliton pulse trains and describe them in a simple way. Consider a train of *g* soliton voltage pulses moving along the *z* axis. The *j* ^*th*^ soliton voltage pulse *V*_*j*_ has a structure of the form, *V*_*j*_(*z, t*) = *e*^−*iπW*^ *V*_*j*_(*a*_*j*_ *z* − *b*_*j*_ *t* + *c* _*j*_), *j* = 1, 2, ‥*g* where (*a*_*j*_, *b*_*j*_, *c* _*j*_) are signal specific pinch deformation parameters. and *W* is the spin structure topology number. We will use this structure in our discussions. We should note that all variables are dimensionless. Thus the position *z* is 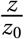, where *z*_0_ is a length scale. Similarly the time variable *t* is 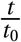 where *t*_0_ is a time scale. Exact analytic results are given in Kalla[23] and Previato[27]. The deformation information carried by transient signals is transferred to create stable long term memories in the pathways traversed by the signals. Let us explain how.

## 4 Memory

### 4.1 Signals align spins to form memory

Information transfer from pulse voltage signals to the pathways they traverse, that have surface spin-half particles on them, follows from the physics laws of electromagnetic theory[28]. Moving soliton pulses carry charge and electro-magnetic theory laws tell us that they produce transient helical magnetic fields around the direction of movement. These transient magnetic fields act on the surface spin half particle, since they are magnetic, and align them to form a non-transient helical aligned spin structure which encodes the transient magnetic field details in it and is thus is a memory. If the structure created is stable it stores the information carried by the magnetic field of the moving soliton pulses as a memory.

The transient magnetic field of a moving soliton voltage pulse has been shown to contains the same information as the voltage pulse that generates it by measuring the helical magnetic field, (Fig 7), produced by an action potential and then reconstructing the action potential[25].

**Figure 7:**
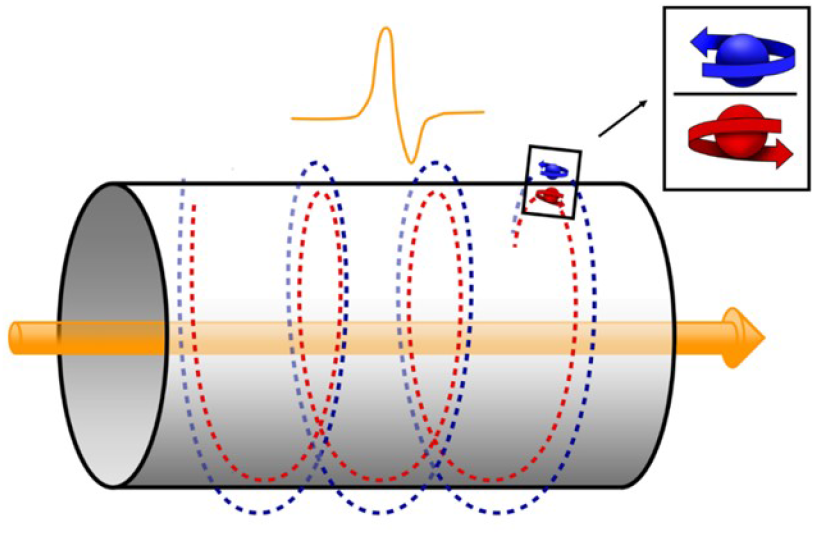
Theoretical helical field generated by a charged voltage puls

Thus the helical non-transient spin structure encodes the details of the transient magnetic field and thus the nature of the action potential that generated it. It is a memory. For memory to be useful it should stable over time and a way of accessing the memory should be in place. We will show that this is possible but for stable memory formation certain conditions have to met by the signals. We will discuss these stability conditions.

We now determine the memory structure, discuss its stability and then estimate its size and the time required to create a stable memory. Our first step is to replace the soliton voltage potentials by an equivalent charge distribution *ρ*, by using Poisson’s equation[28] and then use standard results of electromagnetic theory to determine the magnetic field generated by these moving charges[28]. Poisson’s equation relates voltage to charges by the equation, ∇2*V*_*j*_(*z, t*) = 4*πρ*_*j*_(*z, t*). Using this equation we can replace soliton pulse voltage by their charge density profiles :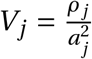, where we have used a simplified representation of soliton voltage profiles that are solutions of the non-linear Schroedinger differential equation. We can simplify the description further by writing, *ρ*_*j*_, as 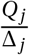 where Δ_*j*_ is an effective width of the pulse and *Q*_*j*_ is the charge carried by the soliton. The speed of the effective soliton charge can now be written down from its profile. It is 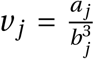 and is signal dependent. Given the equivalent charge profile and its speed we can determine the magnetic field produced by it.

### 4.2 Solitons produce Transient Helical Surface Magnetic fields

The magnitude and direction of the field generated on the surface of a tube of radius *r* by these moving charge pulses can now be written down. The magnetic field has magnitude 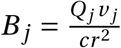, in CGS units, where *c* is the speed of light in the medium, and its direction is tangential to the surface.[28] In terms of ordinates, the field is, 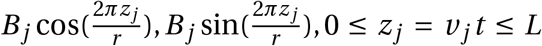, where *L* is a loop length traversed by the pulse. For *N* moving soliton pulses the magnetic field created, 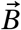, is given by 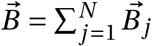. This field has been measured[25] for a single action potential and used to accurately reconstruct it.

Our next step is to work out the alignment of the surface spin-half particles due to this magnetic field. The dynamics of spin-half particles is a governed by quantum theory. We will discuss the details of the structure shortly. For the moment we state that a quantum theory calculation is necessary to study the problem and that the calculation shows that magnetic alignment structure produced is a helical memory structure of spin-half particles. We now identify the nature of the spin-half particles involved and suggested where the structure is located. We next turn to address this issue.

### 4.3 Memory Structures are aligned proton spins

Our network is required to have surface spin-half particles, either electrons or protons, in order to produce solitons. As both of these particles are present on the surface. We now give reasons why they must be protons and we conjecture that the spin memory structure is located in the myelin sheaths of axons. The reason for the choice of protons is that although there are surface electrons in the cations Na+, K+ and Ca 2+ and anions Cl-in the electrolyte. All of these ions have configuration where the 2p orbitals are occupied by two strongly spin-paired electrons. It is difficult o see how these electron spins could freely move to form structures. On the other hand there are unpaired spins of the protons belonging to the hydrogen in water or in the membrane. There are many of them, but their moment is only 1/1800 th of a Bohr magnetron. The small value of the Bohr magnetron makes their interaction very small. Thus their polarisation in the ionic current is even less than that of electrons, but they do have a long coherence time; since the nuclear spn lattice relaxation time may be seconds. The transient helical spin structures might well be be imprinted there.^**^

For this reason we assume the memory structure comes from aligned proton spins. We then show that even though the Bohr magnetron for protons is small the spin interactions are strong enough to form stable memory structures. We conjecture that the protons involved are located in myelin sheaths of axons.

There is some observational support for their conjectured location, which we now discuss.

### 4.4 Are Myelin Sheaths the location of memory structures?

The two fundamental supporting facts for considering proton spins as the building blocks of memory come from MRI measurements. These measurements show that proton spins can be aligned and that protons are present in myelin sheaths. It is also helpful that the relaxation times for MRI aligned protons in myelin sheaths, the T1 relaxation time, is rather long ≈ 10^3^ms. This long relaxation time will allow multiple soliton pulse magnetic fields, each of duration 10^−3^s. to contribute to create the memory structure.

There is more support. Alzheimer, in 1911, had pointed out that the memory loss he observed was due to the degeneration of myelin sheaths. His observation was discarded for many years as the belief was that memories are stored in neurons, but now new research findings support Alzheimer’s original findings[61]. There is now considerable evidence linking memory and myelin sheaths. Here are some facts. It is found that preventing new white matter formation, myelin, hinders learning new motor skills, while structural changes[**?**] in white matter accompany motor training, namely the increase of fractional anisotropy revealed by diffusion tensor imagining methods of MRI after learning. These studies also show that the volume of white matter increases with learning and that the process of myelination is related to the learning of new skills and that myelin sheaths support oscillations[63].

Furthermore, once myelin has formed it is stable with little turnover of oligodendrocytes and limited remodelling of their lengths on existing myelin sheaths. Oligodendrocytes are the cells that myelinate the central nervous system. They are the end product of a cell lineage which has to undergo a complex and precisely timed program of proliferation, migration, differentiation, and myelination to finally produce the insulating sheath of axons. However this stable structure may retain the capacity to remodel if myelin is disturbed[64]. Myelination begins prenatally and continues, in some areas of the brain, into middle age. The time course of myelination varies widely across brain regions. As a general rule, myelination occurs first in neural systems that underlie behaviours that are present early in life. For example, primary sensory and motor areas are myelinated before association areas, and the neural systems involved in postural control and the vestibular sense are fully myelinated before birth. On the other hand, areas of the prefrontal cortex do not become fully myelinated until middle age. By the age of two, myelination is considered to be almost complete, except for the prefrontal cortex[65, 66]. This might explain why our earliest memories begin after that age. It is found the myelin is composed of sheaths, with up to 100 layers for the peripheral nervous system and they are spiral in the central nervous system axons[67]. The presence of multi layers of myelin on an axon, suggests how multiple memories may be stored on a single axon in a time ordered way.

We next move to discuss discuss the memory structure and prove that each structure has a memory specific excitation frequency that may be used to recall the memory.

### 4.5 Mathematical representation of Memory Structures

We first focus on the memory structure and then determine its excitation frequency. Since spin-half dynamics is quantum we have to use quantum methods to study these two features. A quantum calculation [29, 30] (see the Supplementary Information) shows that the helical spin-magnet structure created by the transient pulse magnetic fields of moving solitons, is given by,

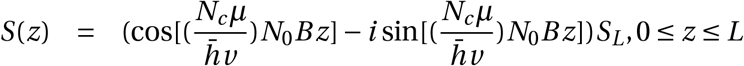

where*N*_0_ is the number of electron in the pulse, *S*_*L*_ is *S*_*i*_(0), the initial spin orientations values at *z* = 0 of the *N*_*c*_ proton spins located on a circle on the axon surface with centre *z*. These spins on a circle on the axon tube are aligned at the same time by the helical magnetic field produced by a soliton pulse. This explains the factor *N*_*c*_ in the expression. The size of a memory will be estimated when we discuss the stability of the structure. There we will estimate the values of *N*_*c*_, *N*_0_

The spin binding is greatly enhanced by topological properties of spins that under 2*π* rotation. Under such a rotation the spin reverses its direction. This leads to a natural pairing of oppositely oriented spins at neighbouring site spins of the helix as they are related by 2*π* rotations. Consequently the memory structure produced has no macroscopic magnetic property. This is an unexpected result.

We next calculate the memory excitation frequency.

### 4.6 Memory excitation frequency

A quantum calculation given in the Supplementary Information section gives a simple formula for the memory excitation frequency. It is,

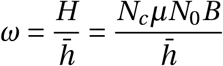

where 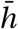 is Planck’s constant, *µ* the magnetic moment of a proton spin, *N*_*c*_ the number of spins on a circle of axon radius at a given point of the axon, and *N*_0_ is the average charge carried by a single pulse. The value of this frequency will be estimated shortly. The range of this frequency depends on the charge carried by a soliton pulse and the profile of the magnetic field and hence is signal dependent. We also show that due to the special topological properties of spin half particles, there will always be a natural secondary memory structure present due to the pairing of two neighbouring spins of opposite orientation located at two neighbouring helical sites. The effective natural frequency of this predicted additional memory system is predicted to have double excitation frequency.

### 4.7 Memory Stability

We now turn to discuss the stability of the memory structure and show that the memory structure of aligned proton spins created by solitons can be stable. The stability argument has three parts. First it is shown that the weak magnetic field of moving soliton’s can align surface spins of protons. Then we show that the the transient magnetic field can create a structure stable. This discussion has several layers. The first layer shows spins can be aligned by the soliton’s magnetic field. In the next layer has two parts it is first shown that one strand of aligned spins is marginally stable followed by a proof of the topological stability of the structure. Topology structure is essential without it the structure can be eliminated by smooth deformations. and possible. At this stage the predicted topologically stable memory structure is in the aligned proton spins on myelin in the axons of pathways between a collection of neurons where topology stability requires the pathways must have loops. This structure looks like an engram. So we call it an engram and estimate its size and the time it takes to form an engram. After these results are in place we show that such a topological memory structure is stable under body temperature thermal fluctuations.

A helpful feature of the structure for stability is that it is held together by the interaction between oppositely aligned spin clusters.^††^

We show that not all signals produce stable memories. We listed three conditions required to create stable long term memories. Let us go through them. The first condition is that the magnetic field *B*_*s*_ must be able to align spins. The second condition is that the memory structure is topologically stable and the third condition is that the structure is stable under body temperature fluctuations.

There are three numbers, *N*_0_ the average charge carried by a pulse, *N*_*c*_ the number of spins influenced by the magnetic field at one point *z* and *N*_*s*_ the number of soliton pulses that decide if the pulses can aligne spins and one additional topological number, the spin topology number *W* is relevant for discussing topological stability of the structure. We first examine the alignment problem.

This requires the aligning energy pulse of duration Δ should be greater than 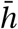. That is 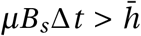. We set Δ*t* ≈ 10^−3^seconds, and *B*_*s*_ = *N*_*s*_ *N*_0_*N*_*c*_ *B*_0_, where the field 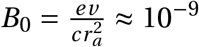 gauss, where *v* is the speed of the signal, *r*_*a*_ the axon radius, the velocity of light, *c* ≈ 10^10^cm/s the speed of light, and *µ* is the proton magnetic moment is *µ* ≈ 10^−23^ in CGS units. For our estimate we have assumed that all pulses produce the same magnetic fields per unit charge, which we set to have the value *B*_0_.

We can estimate *N*_*c*_ it is the number of proton spins in a circle of radius equal to the axon radius *r*_*a*_ thus 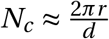 where *d* is the approximate separation between protons. We take *d* ≈ 10^−8^ cm and 2*πr*_*a*_ ≈ 10^−3^cm. Then *N*_*c*_ ≈ 10^5^.

Putting in numbers we get the alignment condition: *N*_*s*_ *N*_0_*N*_*c*_*µB* Δ*t* ≈ *N*_*s*_ *N*_0_10^−30^ > 10^−27^ which gives the condition *N*_*s*_ *N*_0_ >10^3^. Thus a single pulse cannot align spins unless its carries 10^3^ charge units.

We next note that the memory excitation frequency has the theoretical value 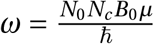. From this it follows that for*N*_0_ = 10, *ω* = 10Hz. If we identify this memory frequencies with sleep spindles oscillations observed, then the value of *N*_0_ is in the range 10 < *N*_0_ < 20. Thus it is predicted that voltage pulses carry very small amounts of charge.

Using this result we conclude that the minimum number of pulses *N*_*s*_ required to align spins is in the range *N*_*s*_ *N*_0_ > 10^3^. We will call this a memory strand. It is not topologically stable.

We next discuss topological stability. Topological stability is essential for the following reason. We would like our results to be relevant for a network that has not just the topological but the real, unknown, geometric connectivity of any brain connectome. This requirement is satisfied by the process of signal generation. Why is this condition important? It is important because if the memory structure is topologically stable it will continue to exist unmodified by topological deformations so that results obtained from our simple topological description of the network remain valid for a network that has the geometric structure of a connectome. Such a smooth transformation exists from the topological classification theorem[32]. A simple condition ensures topological stability. A memory structure is topologically stable only if it has a non zero values for its spin topology number *W*.

### 4.8 Size of an engram

Recall that the spin topology number is defined as, *W* = 4∑ _*i*_ [*α*_*i*_ *β*_*i*_] where *i* ranges over the genus of the signal producing subunit and (*α*_*i*_, *β*_*i*_) are theta function characteristics. The characteristic *α*_*i*_ means a traversal of a loop round the *a*_*i*_ loop and the characteristic *β*_*i*_ means a traversal of a loop round the *b*_*i*_ loop, it follows a memory structure must have aligned spins that traverse both (*a*_*i*_, *b*_*i*_) loops for each value of the label *i*. This is a helical structure. But a set of loops implies the presence of multiple neurons interlinked by axons to form multiple loops in an unknown way. Remember each *b*_*i*_ loop needs to go through two junction points where each junction point is the location of a neuron. Only such an assembly of neurons linked together with loops will have non zero value for the spin topology number *W* and be topologically stable. Thus topological stability requires a structure that describes an engram: a group of linked engram cells with spins arranged in a helical way. This is a theoretical prediction.

The value of *W* for a collection of engram cells has a topological meaning which is easy to identify. It can also be related to the number of neurons *N*_*e*_ in the engram. If all (*α*_*i*_, *β*_*i*_) are non zero for *i* = 1, 2, ‥*k*, then *W* = *k* and *k* is the effective genus of the surface of a specific assembly of engram cells while *N*_*e*_ = *W* − 1. It gives the size of the engram.

A single link of aligned spins, with no loops has *W* = 0, also *W* = 0 if either *α*_*i*_ or *β*_*i*_ is zero. Thus if the aligned spins do not form a helical structure it will have *W* = 0, and the structure is not topologically stable. We suggest that aligned spins with *W* = 0 represent short term memories.

### 4.9 Creation time for engrams

We next derive a formula for the time required to form a *W*≠ 0 memory structure. We showed that a memory link is thermally stable if the memory creating pulse numbers, *N*_*s*_, is greater than *N*_*s*_ > 10^3^, this number assumes that each pulse carries just one unit of electric charge. With this result in place we relate *N*_*s*_ to time *τ*, the time required to produce *N*_*s*_ pulses.

If we call the time required to produce *N*_*s*_ pulses *τ*_*s*_ and the time required to produce one pulse *t*. Then we can write 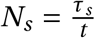. We set *t* ≈ 10^−3^ seconds. Then we have our formula for the creation time of a given memory link is *τ*_*s*_*= N*_*s*_10^−3^ seconds. But topological stability requires multiple memory links that form loops so that the composite memory structure has spin topology number *W* that is non zero. Putting these two results together we have, the time to form a memory engram *τ*_*e*_ is given by,

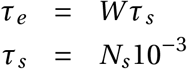

The number *W* is hard to determine but it has a simple topological meaning. It is the effective genus *k* of a specific memory engram. The number*N*_*e*_ of neurons in the engram is related to *W* by the formula *N*_*e*_ = (*W* − 1). Memory structures are interlinked so that removing even a few special neurons can make *W* = 0 and thus affect memory. The time required to form a complex memory formation will involve several independent brain units. For instance for learning juggling eye hand coordination, balance, motor responses are relevant. Each one of these will involve neuron clusters *N*_*i*_ to form the appropriate memory. When activated the number of neurons firing will be 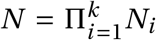, where there are *k* classes of memories that form the engram.

### 4.10 Thermal Stability of Memory Structure

We finally check that our third condition for stability is satisfied. This is the condition that ensures that the aligned spins in a given memory link can withstand thermal disruption when they are bound together by spin-spin interactions.

This requires 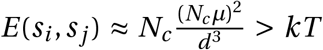. Putting in numbers 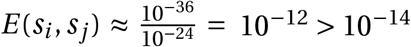. The condition is easily satisfied. The key input was that units of size *N*_*c*_ are naturally aligned at the same time and they thus interact as units.

It is an experimental challenge to either directly verify or rule out the predicted memory structure and the conjecture that it is located in the myelin sheaths of axons. We have provide some indirect evidence in support of the conjecture.

| Variable | Formula(CGS) | Estimate |
| --- | --- | --- |
| Magnetic Field $B_j$ | $B_0 = \frac{ev}{cr^2}$ | $N_1 10^{-9} gauss$ |
| Spin-Field Interaction $H$ | $H = N_s N_c N_0 \mu B$ | $N_s N_0 10^{-30} ergs$ |
| Pulse $H \Delta t > \hbar$ | $N_s N_c N_0 \mu B 10^{-3} > 1$ | Aligned if $N_s N_0 > 10^3$ |
| Frequency $\omega_j$ | $\omega_j = \frac{N_c N_0 \mu B}{\hbar}$ | $N_0 Hz$ |
| Spin-Spin Interaction $S_{ij}$ | $S_{ij} = \frac{N_c \mu_i N_c \mu_j}{d^3}$ | $10^{-12} > kT \approx 10^{-14}$ |
| Spin-Topology $W$ | $W = \sum_{i=1}^k [\alpha_i \beta_i]$ | $W = k, \text{genus of engram}$ |

### 4.11 MRI magnetic fields do not destroy memories

We next show that MRI fields cannot destroy memories. For magnetic fields of a few Tesla strength, the disruption is not strong enough to do this. The disruptive energy is *µB* ≈ 10^−16^ergs is much smaller than the spin binding energy calculated earlier, namely 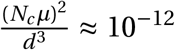 ergs. The binding is between natural units of *N*_*c*_ aligned spins. Thus MRI events can slightly disrupt memories not destroy them.

### 4.12 Table of numbers used to establish memory stability

The stability numbers used are summarized in the table. Values used are *N*_0_ ≈ 10, *N*_*c*_ ≈ 10^5^, *N*_*s*_ > 10^3^, *c* ≈ 10^10^cm/s

### 4.13 Memory Recall: Similarity with Quantum computer

Since memory structures have excitation frequencies, *ω*_*j*_, they could be recalled by a resonance excitation mechanism. The theoretical estimate for memory frequency values found, remarkably, to overlap with those of observed EEG waveforms frequencies. This suggests that memories could be recalled or consolidated by EEG waveforms[33].

The resonance mechanism of memory retrieval has similarities with the way a quantum computer works. In a quantum computer the quantum evolution of a state contains in it all possibilities and the process of identifying a specific possibility is in essence done by a resonance mechanism carried out by the overlap of wave functions, while here the memory space is a collection of different potential memory excitation frequencies encoded in topological stable patterns. An incoming signal with an associated set of EEG frequencies is then a probe which will excite and recall a set of memories by a resonance mechanism.

We next turn to discuss EEG waveforms.

## 5 EEG Waveforms and their Properties

### 5.1 Tiling Waveforms

Topological signal generation deform signal producing subunits of the Riemann surface to topological spheres. We will identify these toplogical spheres as the soma surfaces of neurons. It is expected that these surfaces will have surface voltage oscillations. We will prove the surprising result that the nature of the oscillations is determined by the spin topology number of solitons to be special waveforms that tile the surface of a sphere. This means that identical waveforms allowed, can be placed together without overlapp, Fig(8), to tile the sphere surface. These results are proved using the fact that the oscillations are solutions of the wave equation on the surface of sphere, not a topological sphere. Schwarz in 1873 had shown that tiling solutions on the surface of a sphere exist and that they must belong to five types of waveforms and are related to the existence of the five Platonic solids. Using our dynamic law we will show that the topological sphere surface oscillations can be replaced by sphere surface oscillations and thus the results of Schwarz and the link between soliton spin topology and tiling waveforms that we will prove can be used.

We now identify EEG surface waveforms as due to tiling waveform oscillations on an assembly of neuron surfaces. It is pleasing that their tiling nature is determined by the topological spin carried by solitons. We now show that the surface tiling nature of the waveforms is enough for us to predicts that EEG waveforms will have five frequency bands and determine their allowed frequencies and amplitude values. The prediction is compared them with observations.^‡‡^

Thie suggestion that EEG waveforms are due to the surface oscillations of an assembly of neuron surfaces differs from a standard picture where dipole current oscillations from an assembly of individual neuron dendrites,[2], are regarded as the source of EEG waveforms. The standard approach has nothing to say about EEG waveform frequency bands

### 5.2 Comparison with Experiment

Let us show how the tiling nature of waveforms determines their amplitudes and frequencies. If the waveforms tile the surface of a sphere by identical tiling waveform then each waveform will have a fixed area which is fixed by the number of tiling required to cover the sphere surface. A mathematical result tells us that there are only five such discrete symmetric tiling possible. They correspond to the Platonic solid surfaces. There is one extra infinite class of tiling that can be visualised as orange segment tiling. The number of segments can be arbitrary for this tiling. A simple intuitive argument can now be used to estimate the allowed frequency and amplitude values of the tiling wave forms. The area, *a*_2*n*_, of a tiling solution of a sphere of radius *R* and area *A* = 4*πR*^2^, defines its wave form and is a measure of its amplitude. This result can be understood by considering a small deformation in the radial direction on the surface of a sphere of amplitude *ρ*. The total amplitude *a*_2*n*_ of the deformation is then the sum of all these deformations over the tiling area. It is obtained by integrating the amplitude over one point of the sphere surface over the angles that define the tiling area. Thus, 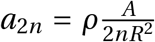, where angular dependence of *ρ* is ignored and*R* is the sphere radius. Now we use the intuitive formula, *a*_2*n*_*ν*_2*n*_ = *c* where *ν*_2*n*_ is the oscillating frequency of the tiling solution and *c* is the speed of the surface wave. Writing 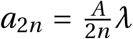, with 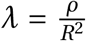, then gives 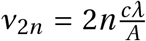, where *A* = 4*πR*^2^, *R* being the radius of the spherical neuron in its excited state. This shows the inverse relationship between the oscillation amplitude and its frequency. A list of our predicted values for frequencies and voltage amplitudes in Table 1 below, is in agreement with observed EEG numbers [44], where the scale is fixed by choosing specific numbers for frequency and amplitude from those allowed for delta wave forms. The amplitude values are in a wide range compared to the frequency band ranges. Frequencies *ν*, in the table, are in Hertz and the voltages are in microvolts. Besides the tiling of the sphere that come from the five regular Platonic solids, mentioned in Table1, there is an additional tiling class for the sphere called the dihedral tiling with orange-segment like tiles.

**Table 1.** The five Platonic solids and general dihedral tiling of a sphere with their predicted amplitudes and frequencies compared with those of EEG brainwaves. Frequencies are scaled by the uniform excitation of the sphere.

| **Solid** | **Tile fraction** | **Frequency** | **EEG Range(Hz)** | **Predicted Amplitude $\mu V$** | **Observed Amplitude $\mu V$** |
| --- | --- | --- | --- | --- | --- |
| Dihedral | $\frac{1}{2n}$ | $\geq 2$ | All waves | $\frac{4\pi}{2n}$ | All waves |
| Sphere | $\frac{1}{1}$ | 1 | delta (.5-3) | 200 | 100 – 200 |
| Tetrahedron | $\frac{1}{4}$ | 4 | theta(4-7) | 50 | < 30 |
| Cube | $\frac{1}{6}$ | 6 | alpha(8-13) | 33 | 30 – 50 |
| Octahedron | $\frac{1}{8}$ | 8 | alpha(8-13) | 25 | 30 – 50 |
| Dodecahedron | $\frac{1}{12}$ | 12 | beta(14-30) | 17 | 5 – 20 |
| Icosahedron | $\frac{1}{20}$ | 20 | beta(14-30) | 10 | 5 – 20 |
| Dihedral | $\frac{1}{40}$ | 40 | gamma(31-50) | 5V | 5 – 10 |

The area value of a given tile can be exactly determined by using a result of spherical geometry. If (*α, β, γ*) are the angles of a spherical triangle for a sphere of radius *R*, then its area *A* is given by the formula 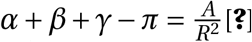. If these angles are chosen to be 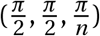 then its area is 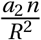, where two of these triangular sections are used to represent a wedge. These wedge tiles the sphere surface into 2*n* pieces as stated earlier, with frequencies proportional to 2*n*. An analytic calculation[45] shows that the frequency is 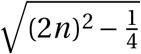. Thus it supports the simple relationship proposed. The frequency numbers depend on the choice of scales. These are fixed by setting the frequency value of the delta wave form to be one and its amplitude 200*µV*. We note that dihedral tiling (orange-segments) can produce all observed frequencies. An illustrative example is given that produces a specific gamma frequency. The analytic calculation[45], also give explicit expression for the tiling solutions that demonstrate their local nature, namely that a given oscillation is confined to one tiling segment.

The analytic way to determine the nature of sphere surface oscillations is to solve the sphere surface wave equation and impose suitable boundary conditions to produce tiling waveforms. This step requires replacing a topological sphere by a geometric sphere which can be justified, as we will show, by using the dynamical principle of the network.

We now have a surprise. We find that the boundary conditions required to produce tiling waveforms are automatically fixed by the input soliton spin topology phases that enter a neuron to produce a signal. This is an unexpected result. It directly relates EEG waveform properties to topological properties of brain excitations. Let us prove this result.

### 5.3 Tiling waveforms are solutions of a Wave equation

Recall a wave equation on any surface has the structure,

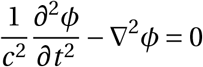

where ∇^2^ is the Laplacian operator appropriate for the surface. The dynamical law requires all valid results of the surface must respect its underlying Riemann surface structure. The topological sphere is a Riemann surfaces that is conformal equivalent to a sphere [37]. Conformal symmetry is a basic property of any Riemann surface[20]. Thus the dynamical law requirement that only results that respect the underlying Riemann space structure are valid means that only those results that are invariant under conformal transformations are valid. Since a topological sphere is conformal equivalent to a sphere we solve the wave equation on the sphere and determine the allowed frequencies and waveforms. The oscillation frequencies found are invariant under conformal maps, but the Laplacian operator and waveforms shapes are not, it follows that our results for frequencies are in agreement with the dynamical principle and are valid results but waveform shapes obtained are valid to within a conformal transformation.

We need one more mathematical detail. The Laplacian operator must have the symmetry of the sphere surface. This leads to the result that radial part of the wave equation is the linear second order hypergeometric differential equation[38], on the surface of a sphere, with three regular singular points[36]. We identify these three singular points as the location of three primary dendrites on a neuron surface. The median number of primary dendrites has been measured and found to be between three and four[39, 40] and the range of primary dendrites observed varies between two to seven. We will consider the case of three dendrites. The analysis for the case when there are more singular points is possible[36].

### 5.4 Tiling waveforms are fixed by spin topology phase

The spin topology number phase is given by *e*^*iπW*^, 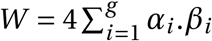 and (*α, β, i* = 1, 2, ‥*g*)[2] are the discrete characteristics of the Riemann theta function, associated with the the two classes of topological loop coordinates (*a*_*i*_, *b*_*i*_) of a Riemann surface (Fig 3) where each characteristic (*α,β*_*i*_) can only take one of two values, namely 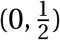,. It is a global property of the subunit and depends on all the loops of the subunit that generates a signal. The topological number has a simple geometrical meaning. It tells us the pathways involved in producing the soliton since the characteristic*β*_*i*_ associated a loop *b*_*i*_ while the characteristic *α*_*i*_ is associated with a loop *a*_*i*_ of the Riemann surface (Fig 2). The two characteristics thus define helical loops along the (*a*_*i*_, *b*_*i*_) loops of the Riemann surface. Notice that for *W* to be non zero the memory path should be helical.

Let us next explain why phases at singular points determine the nature of solutions. Suppose *z* = 0 is a regular singular point and the solution of the hypergeometric equation looks like *Az*^*µ*^ near *z* ≈ 0, where *A* is a constant and *µ* is not an integer. It is called the index of the singular point. If we replace *z* by *ze*^2*iπ*^ the solution changes from *Az*^*µ*^ to *Az*^*µ*^*e*^2*iπµ*^. If *µ* was an integer the factor *e*^2*iπµ*^ = 1 but when it is not we have a phase which describes the nature of the singular point. A mathematical result[36] tells us once *µ*, is fixed the nature of the corresponding solution of the hypergeometric differential equation[3] is determined.

We will now related these singular point phases to the input soliton’s spin topology phases *W*_*i*_, *i* = 1, 2, 3. Let us follow the procedure of Schawrz. The hypergeoemtric equation is a second order differential equation and thus has two linearly independent solutions. Schwartz considered the ratio of the two linearly independent solutions that had the property that vanish at each of its the three regular singular points. The three singular points as the vertices of a triangle on the surface of a sphere. The vertices form a spherical triangle.

Then solutions were constructed that vanish along the edges of the spherical triangle and showed that the angles of the triangles were given by the difference of the indices of the three singular points. Thus if the angles, related to the parameters of the hypergeoemetric equation, were chosen appropriately the solution waveform would be a localised oscillation within a region and would have the correct area to tile the sphere surface. Recall for a spherical triangle its angles fix its area. Let us write down these angles for the hypergeometric equation‥

The hypergeometric differential equation is given by[3]

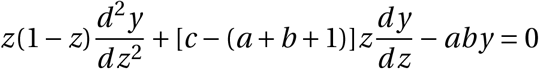

where (*c, c* − *a* − *b, a* − *b*) are real but not integers. The three singular points are (0, 1, ∞) where ∞ is the north pole of the sphere, and the points (1, 0) are on the equator. For such a triangle the angles of the solution triangle described, were found by Schwartz to be *π*|1−*c*|, *π*|*c* −*a* −*b*|, *π*|*a* −*b*| corresponding to the points (0, 1, ∞).

Schwarz set 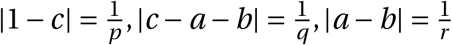 where (*p, q, r*) were integer greater than or equal to two and showed that a spherical triangle with angles 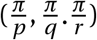 tile the sphere provided 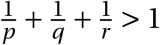. There are only four possibilities: dihedral tilings where (*p* = 2, *q* = 2, *r* = 2*n*) and *n* = 2, 3, … The other tiles correspond to tetrahedral, octahedral and icosahedral tiles As Yoshida[36] shows these tiles can be related to the surfaces of Platonic solids and the solutions are determined by these angles. We now show that input soliton spin topology phases entering at the three singular point locations, identified as dendrite locations, fix these angles.

Let us now write the angles of the solution triangles as phases 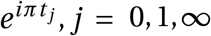 if 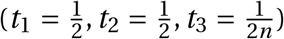 where *n* = 1, 2, ‥ then these angle choices produce a dihedral tiling of the sphere. The corresponding phases are those associated with a solution of the hypergeometric equation near that singular points. We will call the phase *e*^*iπt*^ a twist angles.

These twist angles are fixed by the soliton spin topology phases: it is a dynamical mechanism. We assume that a unit twist *W =* 1 associated with a soliton spin phase produces a twist of, *e*^*iπt*^ to a solution at a singular point of the hypergeometric differential equation. Then a topological twist *W* produces a twist (*e*^*iπt*^)^*W*^ = *e*^*itπW*^. Next we note that a set of characteristics is said to be an even/odd depending on whether *e*^*iπW*^ = ±1[20]. Solitons can have either odd or even characteristics so that the constraint of the topology phase they carry is simply *e*^*iπW*^ = ±1. For a given soliton *W* can be either an even or odd integer.

Thus we require *e*^*itπW*^ = ±1, that is 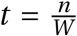 where for a given soliton 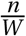 is either an even or an odd integer. For example set of values of *t* for the three regular singular points can be 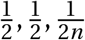. This corresponds to the dihedral tiling example, we used, and corresponds to *W*_1_ = 2,*W*_2_ = 2,*W*_3_ = 2*n*. The technical details are in Yoshida[36].

Fifteen explicit tiling soltions of the linear hypergeometric differential equation were discovered Schwarz[35]. Later a more comprehensive list of such solutions was found[42, 43]. An example of a tiling waveform is shown in fig 4.

### 5.5 Special Tiling Waveforms: Fractional Legendre functions

We now depart from discussing general tiling waveforms and restrict ourselves to the the dihedral tiling. The advantage of doing this is two fold. First, explicit analytic expressions for these tiling solutions and their frequencies are known and second these solutions form a complete set. This means that any arbitrary function on a sphere can be expressed as a linear combination of these tiling solutions. They can thus be used to represent all other tiling solutions.

The analytic expressions for dihedral, cubic and tetrahedral tiling are known. They are fractional Legendre functions,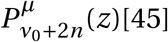. These tiling solutions form a complete set so that they can be used construct Greens functions *G*(*z, t* : *z*^′^, *t* ^′^)[3] with the help of Greens functions we can determine the output waveforms *H* (*z, t*) that will be produced in response to a given input *s*(*z*^′^, *t* ^′^)‥ The mathematical details of the construction and the form of *G*(*z, t* : *z*^′^, *t* ^′^) are given in the Supplementary Information(SI). The completeness property also allows us to write any function *f* (*z*) on the sphere in terms of the fractional Legendre functions as,

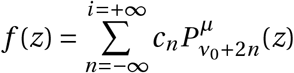

where the points *z* represent points on arcs on the sphere. A slight generalisation allows us represent not just functions on arc of the sphere but arbitrary functions on the sphere surface by considering the analogues of the usual spherical harmonics[3] for fractional Legendre functions. The ones related to dihedral tiling correspond to setting 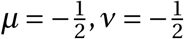. They are:

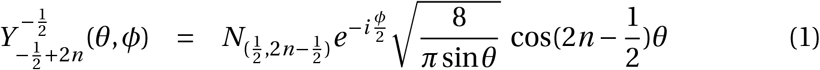

where *n* = 1, 2, ‥ and 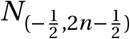 is a normalization factor. They are evaluated in the next section. These functions are plotted for special values of *N* (Fig 7,8) to show the two dimensional tiling nature of these functions and that the solutions are localised. These functions also form a complete basis set of functions so that now any arbitrary function on the sphere surface can be represented using them.

**Figure 8:**
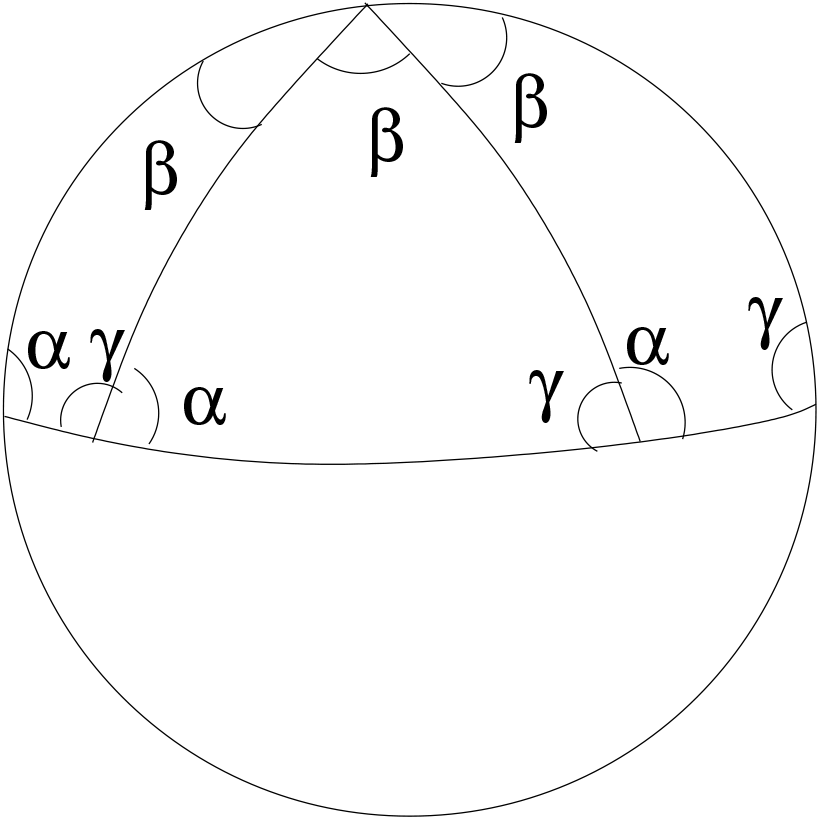
Schwarz Tiling of a sphere with dihedral tiles of equal area. The spherical triangle with apex angle and base angles 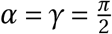 and sides that are great circles tile the sphere if the apex angle 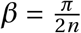 where *n* is an integer. Possible tiling are related to Platonic solid surfaces projected onto the sphere. For example, an octahedron with eight triangular faces has *n* = 4 and 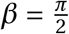, giving a tiling with eight triangular tiles, like the one illustrated.

### 5.6 Plots of EEG waveforms in the dihedral approximation

A representative plots for the theoretically determined delta and theta waveforms are displayed using Eq(7) in Fig(9) and Fig(10).

**Figure 9:**
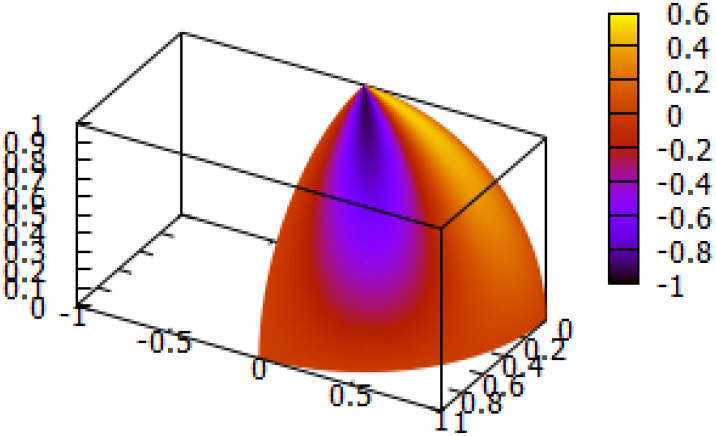
Theoretical Tiling Delta Waveform using Eq(7), n=1. The figure shows how an assembly of neuron surface delta oscillations will appear in the brain, represented as a hemisphere of radius one. Note the oscillations are localised to a tiling sector of the hemisphere. The colour coding show that the boundaries have zero amplitude

**Figure 10:**
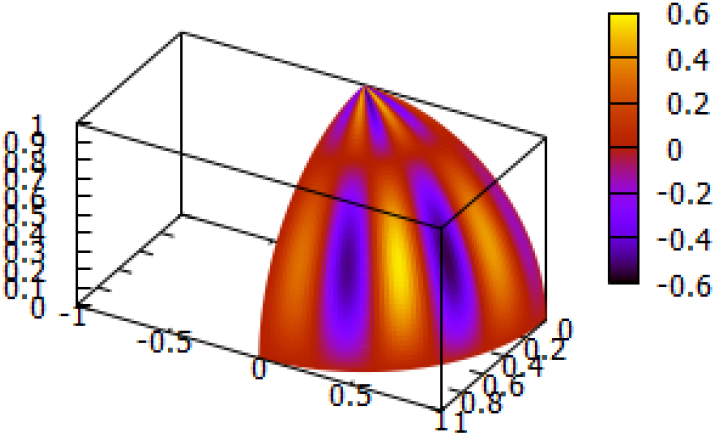
Theoretical Tiling Theta Waveform using Eq(7), n=2. A second example. Note the tiling property.

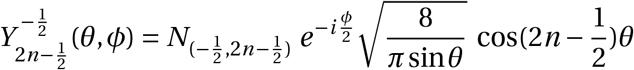

where *n* = 1, 2, ‥. The angles (*θ, ϕ*) are spherical polar coordinate angles with ranges (0 ≤ *θ* ≤ *π*, −*π* ≤ *ϕ* ≤ +*π*).

To construct tiling solutions that vanish on the sides of a spherical triangle the following variable changes are required 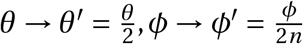 and we require that the solution be periodic in *ϕ*^′^. In terms of these variables, the solution plotted are,

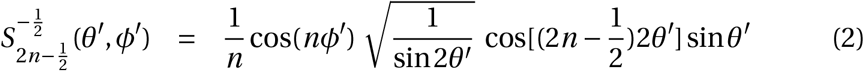

where the range of the angles are now 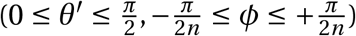. We check that this solution vanishes when 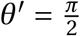 and when 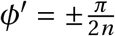 which define the three sides of a spherical triangle. Thus we have constructed a tiling solution.

### 5.7 Prediction of EEG waves generated by input signals

The special Greens function constructed in the Supplementary Information, can be used to determine the distribution of dihedral tiling EEG waveforms *m*(*z, t*) generated in response to any input signal *s*(*t, z*) by the formula,

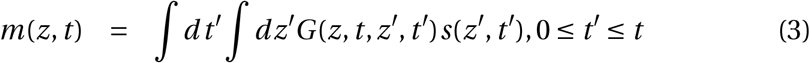

Consider two examples of input signals. Both are local signals, where one is a transient signal and the other an oscillatory signal. We have, 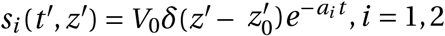 with *a*_1_ = *a*_0_ > 0, *a*_2_ = *iω*_0_. In the case of a transient input signal, delta waves and other oscillatory waveforms are produced, while in the case, of an oscillatory input, if its frequency *ω*_0_ overlaps with any EEG waveform frequency *ω*_*n*_ then the the predominant excitation output will be EEG waveform of that frequency. This is an important remark. It tells us that transient high energy input signals can produce slow oscillatory EEG waveforms.

The analytic solutions for the dihedral tiling waveforms, labelled by an index *n*, allow us to write down an analytic formula for their allowed frequencies[45]. We have,

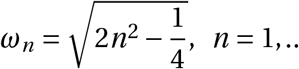

however, the scale of these frequencies is not fixed by the equation. We have set the scale to be one. In Supplementary Information we use an energy argument which justifies this scale choice. The waveform shapes of these dihedral tiling solutions are given by the fractional spherical harmonics. They represent 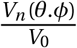, the voltage of the waveform where *V*_0_ is a scale to be chosen from experiment. Thus

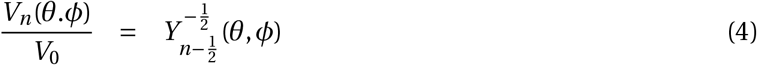

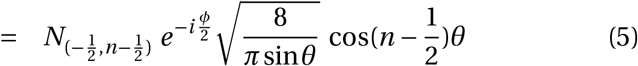

where *n* = 1, 2, ‥ and the real part of the harmonic is to be used. Thus we have explicit expressions for surface waveforms and a formula for their allowed frequencies. The normalization factor 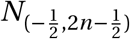 can be calculated by using the Maier’s[45] results relating 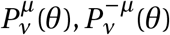. Their norm 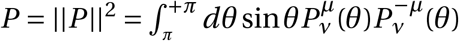 can now be evaluated. We have,

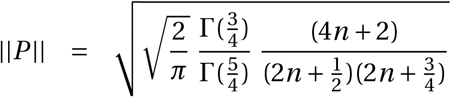

This result is required to calculate Green’s functions.

Two practical uses of EEG waveforms are given in the Supplementary Information sections. They show how how to represent an arbitrary surface function using the dihedral waveforms and how the location of a brain excitation can be unravelled from EEG waveforms.

However these theoretical results simply use the mathematical idea of completeness and have limitations. Any input signal to the brain initiates brain processes but since the brain is an open system, besides the input signal introduced, other internal or external signals will be entering the brain that will have consequences. These unknown deviations due to the open nature of the brain cannot be included in the theoretical formula. Consequently the predicted mix of EEG waveforms for a given input will have deviations. In our view these deviations provide insights regarding brain processes.

We next consider a sequence of EEG modulations observed during deep sleep and illustrate how the observed events can be understood as brain processes by the use the properties of EEG derived. We then outline the way neural field model these events to highlight the differences between the two approaches.

## 6 EEG modulations observed in deep sleep

We consider a sequence of EEG modulations observed during deep sleep, the K-complex modulation, followed by sleep spindles, followed by sharp wave ripples[46]. We interpret these events as brain processes that consolidate and transfer memory. The account we give suggest that EEG waveforms play an important role in memory consolidation and transfer.

Sleep occurs in five stages: wake, N1, N2, N3, and rapid eye movement(REM) sleep. Stages N1 to N3 are non-rapid eye movement (NREM) sleep, with each stage leading to progressively deeper sleep. Approximately 75% of sleep is spent in the NREM stages, with the majority spent in the N2 stage. A typical night’s sleep consists of 4 to 5 sleep cycles, with the progression of sleep stages in the following order: N1, N2, N3, N2, REM. A complete sleep cycle takes roughly 90 to 110 minutes. The first REM period is short, and as the night progresses, longer periods of REM and decreased time in deep sleep (NREM) occur.[47] The K-complex modulation appears in the N2 sleep phase, and is followed by sleep spindles.

We interpret the observations as a sequence of brain events that are initiated by a major blocking excitation that leads to the K complex modulation and follow through the expected consequences of this major modulation and show this suggested chain of expected brain events explains sleep spindles and sharp wave ripples. We also make a general prediction. Signal blocking events, we believe, are essential for the functioning of the brain as they allow focused attention. Consequently we expect such events to happen constantly at different energy scales and will show up in different EEG waveforms as sharp spikes[56]. Understanding the process of how they are triggered is a challenge.

### 6.1 Facts about K-complexes

In the N2 sleep cycle, delta waves are observed and, at intervals of 1.0-1.7 minutes large sharp spikes, known as K-complexes, appear with amplitudes in excess of 100 mV. They modulate high amplitude delta waves (1.6-4.0) Hz. The excitations have duration greater than 500 ms and are followed by spindle shape oscillations of delta waves, of duration a few seconds. The modulations are in the frequency range (10 − 15) Hz, and can be followed by sharp wave ripples [50, 51, 52]. It is likely that the high voltage low frequency waves are not delta waves but EEG waveforms generated by major blocking events that occurs during deep sleep, since signal blocking excitations are known to result in EEG waveform production[53]. These waveforms, are Slow Oscillating waves. We call will them delta waveforms because all surface oscillations observed, in the network, come from neuron surface oscillations.

The K-complex sequence can be modelled in our approach in the following way: a major blocking event produces a modulation of a high voltage delta wave which is then modulated by the K-complex event. The high voltage delta waveform, then excites an underlying memory structure and its excitation frequency modulation of the delta waveform produce the sleep spindle modulations, which then, excites an underlying memory by a resonance mechanism that shows up on a delta wave as sharp wave ripples[49]. This sequence events is an example of a feedback loop of information exchanges between signals mediated by EEG waveforms. It can be interpreted as a memory transfer from one brain location to another.

### 6.2 Transient Localized Signals generate K-complex

Let us now give the mathematical details. Explicit analytic expressions for transient localised solutions of the non-linear Schroedinger equation, that can cause signal bocking, exist[23]. To simplify our analysis we approximate the known analytic solutions by the following simple expression,

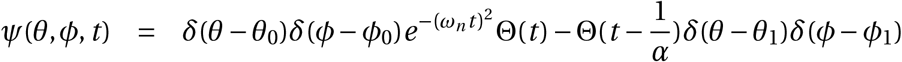

where (*θ*_1_ = *θ*_0_+Δ, *ϕ*_1_ = *ϕ*_0_+Δ), and Θ(*t*) = 1, *t* > 0 and zero otherwise, the Heaviside Theta function. For our qualitative discussion we drop normalization factors and ignore the spatial oscillations of the waveforms. The theoretical surface waveform modulation due to a transient localized excitation is obtained by simply superposing this transient excitation voltage form with the delta waveform voltage. This is plotted in (Fig 11) using,

**Figure 11:**
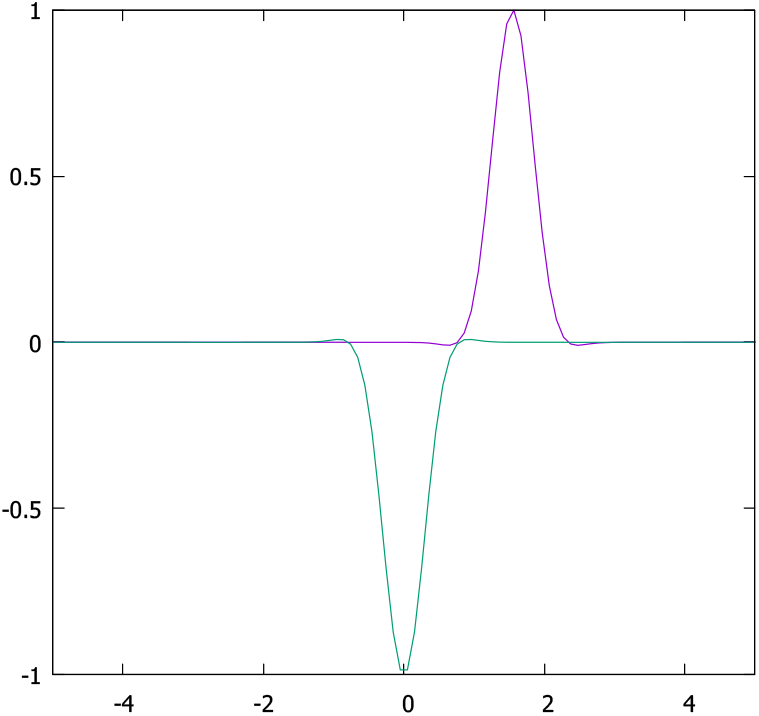
Theoretical K Complex like Structure using Eq(5),Eq(6),Eq(7). Scales are arbitrary

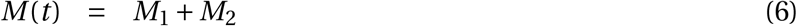

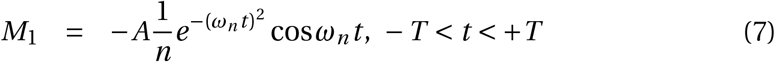

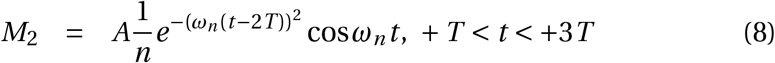

with 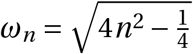. For numerical work an appropriate gluing function *M*_12_ is required and a value of the constant *A*, has to be chosen. This is the K-complex. It is a modulation of a delta waveform by a transient localised blocking excitation.

### 6.3 Modelling Sleep Spindles

We next suggest that sleep spindles are induced modulations of delta waveforms due to the excitation of helical magnetic memory structures discussed earlier. The modulation occurs in two step process.

In the first step the surface charges of the memory structure are acted on by the EEG waveform voltage. As the delta waveform voltage is high this excites the memory to its natural frequency. This induced memory oscillation produces a potential Δ*V* that then modulates the EEG potential to produce a sleep spindle. Sleep spindles are thus interpreted as copies of memory structure frequencies.

We showed earlier,Eq(4), that the delta waveform potential, in the dihedral approximation, was,

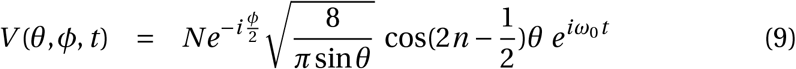

where the time dependence of the oscillation is added and the voltage scale is included in the normalisation constant *N*. The spherical polar coordinates, have constraints, 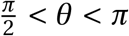, and 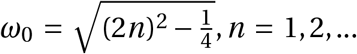 The gradient of −∇*V* is an electric field that acts on charges *e* on the surface causing them to move. We replace 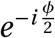 by sin(*ϕ*) and suitably adjust its range.

We next replace the (*θ, ϕ*) dependence of *V* (*θ, ϕ, t*) by a simpler expression but we retain its theoretically determined time dependence. Thus we set *V* = *V*_0_ sin *θ* sin *ϕ* sin *t* for the delta waveform, so that *ω*_1_ ≈ 1. Then *V* (*t*) = *V* (*x, y, t*) = *V* (*x, y*, 0) sin *t* in Cartesian coordinates. Thus we have electric fields induced by the delta waveform potential gradient in the x and y directions that acts on an electron of momentum 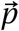. We have,

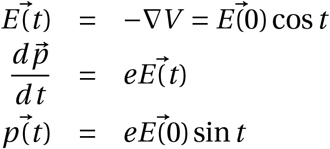

where 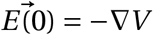. This interaction causes a displacement of the electron and gives it energy. If this energy is high enough it excites the memory structure making it oscillate with its natural frequency 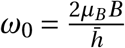 and an additional oscillation of frequency 2*ω*_0_ that is due to the paired spin structures discussed earlier is also excited. To excite the memory structure we require that *eE* (0).*ds* > *NE*_*M*_(0) where *eE* (0).*ds* is the electron energy correponding to a displacement of the electron by *ds*, the memory structure energy is 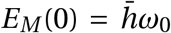, and *N* is the number of electrons in the memory structure. Once this threshold is reached the memory structure is excited and each electron oscillates with energy 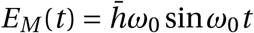. We identify this oscillating energy term as the modulation voltage, *e*Δ*V* seen as a sleep spindle. We have,

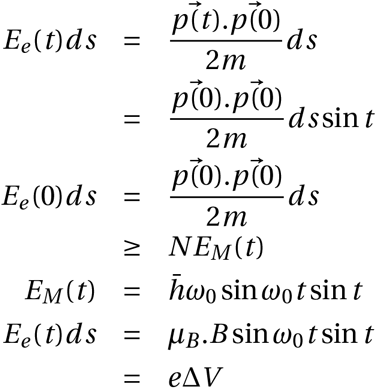

Thus the modulation due to a single electron is given by 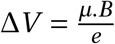 sin *t* sin(*ω*_0_*t*), 0 ≤ *t* ≤ *T*, while the observed modulation involves the cluster *N*_*s*_ ≈ 10^3^ that we found were necessary to form memories. We have used the result 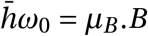. The modulation due to the original spin memory structure is thus,

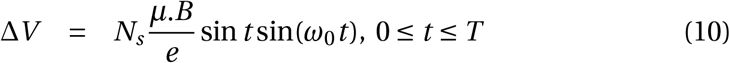

There is a corresponding Δ_2_*V* modulation due to the induced electron spin paired memory structure, given by 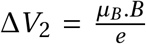 sin *t* sin(2*ω*_0_*t*)*ds*, 0 ≤ *t* ≤ *T*, where *ω* is the original memory structure frequency. The oscillation frequency of the memory has been written as sin(*ωt*). The scale of the sleep spindle figure (Fig 12) shown is arbitrary and we have only plotted Δ*V*_1_.

**Figure 12:**
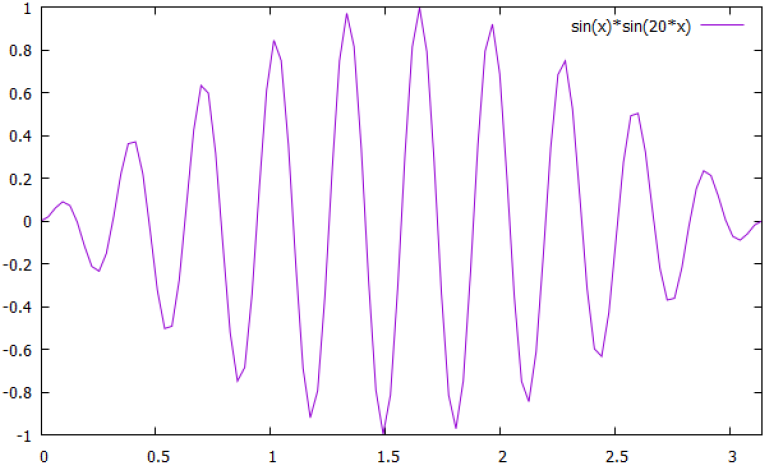
Theoretically determined Sleep Spindle Structure with only the original memory structure considered Eq(9). A second memory structure with double frequency and double amplitude is theoretically predicted. See text for details.

Sleep spindles have observed frequencies in the range *ω*_0_ ≈ 10−15 Hz which is consistent with the theoretical estimate.

### 6.4 Interpretation of deep sleep events

The sequence of EEG modulations observed is interpreted as a mechanism of memory consolidation and memory transference from one region of the brain to another [55, 70, 71]. Major blocking event can creates delta waveforms[53]. The network provides theoretical justification for such an interpretation. Our discussion of sharp wave ripples did not include details regarding them or provide an analysis of what inferences can be drawn from their observed features. We now provide these details. We then show how using our results regarding EEG generation and interpreting the sharp wave ripple as a modulation caused by a memory transference signals lead to the estimate the number neurons firing that accompany it‥

### 6.5 Comments on Sharp Wave ripples

Let us state some facts about Sharp wave ripples. They modulate delta waveforms by frequencies in the ranges, (80−1500)Hz, and also in the ranges (20−40) Hz and by one of frequency ≈ 5 Hz. Sharp wave ripples are a major brain event that can involve between 50, 000 to 100, 000 neurons firing[48]. They follow sleep spindle events.

From the network perspective a resonance induced signal, like all brain excitations, is topologically produced and thus can be described by a pinch deformation. This means EEG waveforms will be co-produced. The nature of these EEG waveforms, as we showed, is determined by the unknown spin topology numbers *w* of the generated signals. We recall how this works using a dihedral tiling to illustrate the process. Recall that three input spin topology numbers *w*_*i*_, *i* = 1, 2, 3) entering the three primary dendrites of a neuron fix the nature of oscillations on the neuron surface. For dihedral tiling these three input numbers are (2, 2, 2*n*) and they produce an EEG waveform of oscillating frequency *ω* = 2*n*. Thus by observing EEG oscillations the spin topology number *w* of a signal can be determined.

The value of *w* also, as was pointed out, reflects the size of a memory circuit, its effective genus *k* which in turn is related to the number of neurons *N*_*k*_ that define the circuit. We have *w* = *k* and *N*_*k*_ = (*w* −1) Thus to create frequencies in the range (80 −150) we need (80 < *w* < 150). Similarly to produce the frequency range (20−40) we need spin topology numbers in the range 20 < *w* < 40 and for ≈ 5 a *w* ≈ 5.

We now assume that the observed EEG waveform modulations in a sharp wave ripple t are related to memory engram neurons firing. This will involve engram neurons for the different sensory memories. From the observed sharp wave ripple EEG waveforms we can extract the following memory engram numbers : (*N*_*v*_ ≈ 10^2^, *N*_*a*_ ≈ 10^2^, *N*_*t*_ ≈ 10, *N*_*s*_ ≈ 5), where *N*_*v*_ is the vision engram size, *N*_*a*_ the audio engram size,*N*_*t*_ is the touch engram size and *N*_*s*_ is the smell engram size. Multiply these numbers we get an estimate of the expected number *N*_*E*_ of firing neurons. We get *N*_*E*_ ≈ 50, 000.

This sketch suggest how observational data can be interpreted using the networks vocabulary. Thus for sharp wave ripples, we used the EEG frequencies observed to predict the expected number of firing neurons. We have related the high frequency EEG components with vision and audio senses, the lower frequency EEG waveforms touch and smell.

Our prediction is that any structure supporting memory should exhibit the frequency ranges similar to those found for sharp wave ripples as they reflect memory recall and transfer processes. The memory structure is in the pathways but the excitation frequencies are triggered by the firing of neurons that are linked together by these pathways to form an engram. The associated EEG waveforms are co-produced during this process contain in them the firing neuron details that create them.

We now briefly comment on the difference of the network approach to model brain events with that of neural field theory by examining how neural field methods model the K-complex sequence. In the network approach, complex processes of the brain are related to a sequence of elementary processes that involve the interplay of brain signals, EEG waveforms and memory structure using biology and physics inputs. The inclusion of feedback loops of interactions between memory and signals is a key process. In our discussion of K-complex events this approach was used. This was possible because the network suggests how EEG waveforms are related to other brain signals and suggests how and where memories are stored and suggests a way EEG waveforms can interact with memories.

In neural field theory all brain processes involve a competition between inhibitory and excitatory neurons that is process dependent. This powerful insight coupled with biology driven interactions allows neural field theory to model a variety of brain processes. The focus in neural field theory when modelling the K-complex sequence of events is to selectively understand specific excitations observed, such as Slow Wave Oscillations and K-complex[52], sleep spindles [57, 58] and memory transfer[59]. There is also difficulty in formulating the problem[52]as K-complex events are modulations on an elevated voltage background. EEG waveforms are introduced as induced excitations due to dipole current loops on dendrites. Such a model does not produce the observed frequency amplitudes of EEG waveforms and does not suggest how interactions between signals and memory occur. Finally in modelling memory transfer the neural field theory approach does not relate memory transfer to observable events such as sharp wave ripples but to long term age related degeneration of memory.

## 7 Discussions

In this paper a different topological way of thinking about brain signals, how they are created and how memories are formed is suggested. Signals were generated by pinch deformations of axon tubes. This idea has biological support[60]. Pinch deformations are the input signals of the network and can be created by electric, mechanical or chemical means. The mathematical parameters that describe pinch deformations come from the Riemann structure of the surface network. Thus the process of signal generation autonomous. We also explained the way memory structures of aligned proton spins could form and be recalled by their memory specific excitation frequencies.

### 7.1 Fundamental Brain Processes

The central idea is that global topological features of the brain may provide insights that are otherwise not available. To implement this global approach we used mathematical ideas of topology, algebraic geometry and physics. However even though the mathematical ideas used are unfamiliar, we emphasise that the new approach offers an intuitive way of thinking about brain processes. The approach suggests that all brain processes involve a small number of elementary sub processes such as the blocking signals, resonance interactions between EEG waveforms and memory, modulation of waves, and induced feedback loops between signals and memory. In this list understanding blocking signals, how they may be generated by a person by chemical means is a key feature. Blocking signals isolate a system so that it can focus on specific events. It is an essential feature of all cognitive functions of the brain. Each subprocess has an intuitive meaning. We showed how interactions between memory and signals could be described and intuitively understood.

We next summarise some of the key results established.

### 7.2 EEG Creation

The suggested method of generating EEG waveforms as special surface voltage oscillations on spheres created by pinch deformations immediately links them to these pinch deformation created excitations and show that they tile the surface of a sphere. From this we could prediction their frequencies and amplitudes. Surprisingly the frequency band frequencies of EEG waveforms overlap with the memory excitation frequencies theoretically predicted. This surprising because the origin of the two frequencies are very different, one is a classical system describing surface oscillations while the other is a result for a quantum system acted on by a magnetic field. This overlap of frequencies, however, plays an essential role in the functioning of the brain as it allows EEG waveforms, co produced with an incoming signal, to excite memories by the mechanism of resonance excitation and thus identify its nature.

Two further practical results are derived, and placed in the Supplementary Information section, because of their technical nature. One predicts the mix of EEG waveforms expected in response to an arbitrary input voltage signal and the other reveals the EEG structure of an observed brain excitation event. The first result is expected to have deviations for two reasons, first new brain events occurring during the period of observation and second if the input signal initiates additional brain excitations that are not represented in the mathematical scheme. Both these deviations thus throw light on the way the brain functions and make the procedure is a useful probe. The second reveals the extent of the event, that is, the number of neurons involved in its creation.

A major result of the approach was to predict a memory structure that had structural features similar to engrams but with the additional prediction that the memories were in the pathways between neurons.

### 7.3 Engram location and properties

Memory structures naturally emerge in the surface network. The first step is that pinch deformations generate signals that carry information are charged and by the laws of physics produce magnetic fields that in turn act on surface spin-half particles to create topological memory structures with memory specific excitation frequency. We conjectured that the memories are stored in helical aligned proton spins with non zero spin topology number *W*, located in myelin sheaths of axons.

A non zero spin number means that the structure must include loops of aligned spins but loops in the network have neurons at their junction regions. Thus the theoretical topological stable memory structure predicted is one of a group of neurons linked together by pathways with non trivial topology, where the memories are in the pathways. This predicted structure has the features of observed memory engram.

We derived a formula for the creation time *τ* for an engram and its size in terms of the spin topology number *W*. An engram can encode multiple memories. Each memory engram can encode a specific subset of memories and will have its own non zero *W*_*e*_ value. The creation time *τ* for such an engram we showed is given by the formula,

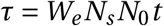

where *N*_*s*_ is the number of pulses required to create a thermally stable memory strand, *N*_0_ is the average charge carried by a pulse and *t* is the time required to generate one pulse. We found the condition for aligning spins was *N*_*s*_ *N*_0_ > 10^3^. By identifying sleep spindle frequencies with memory excitation frequency we found *N*_0_ ≈ 10. Thus we choose *N*_*c*_ ≈ 10^3^ as the number of pulses that can stably align spins. Finally we ser *t* ≈ 10^−3^, seconds as a measure of the time taken to create one pulse. Our memory creation time then becomes, *τ* ≈ *W*_*e*_ seconds. The spin topology number *W*_*e*_ can be related to the number of neurons *N*_*e*_ in the memory unit, its size, by the formula *N*_*e*_ = (*W*_*e*_ − 1). But this time estimate does not include the additional time required to myelinate axons, if necessary for storing the new memories.

Aligned spins with *W* = 0 that do not form loops are not topologically stable memories. We interpret them as short term memories. Their lifetime is expected to be related to the relaxation time T1 of MRI for white matter.

We conjectured that the memory structure is located in myelin sheaths of axons. We offered some evidence that this may be the case. We now make an additional prediction in support of the conjecture. If myelin sheaths do store memories then they should involved in producing sharp wave ripples. Recall, we interpreted sharp wave ripples as a signature of a memory transfer event. Results established for the networks predict that the nature of observed EEG frequencies is determined by the three spin topology numbers *w*_*i*_ .*i* = 1, 3 of signals entering a neuron through its three primary dendrites. The value of *w*_*i*_ depends on the number of neurons *n*_*e*_ involved in the process, and is given by (*n*_*e*_ = *w* − 1). For dihedral tiling the three spin topology numbers are (2, 2, 2*n*) and they produce an EEG frequency *ω* = 2*n*. These frequency for the sharp wave ripple were (80 − 150, 20 − 40, 5) Hz. These EEG frequencies values are expected to be present in all memory structure of the brain and hence should be present in memory recall and transfer processes involving engram neurons.

### 7.4 Brain Excitations can be chaotic

In our discussions all excitations considered were non chaotic[11]. However chaotic brain excitations can be produced in the system in two ways. The first is by the coupling of two oscillatory modes of different frequencies, for example two large brain excitation such as a K complex, close together can couple EEG waveforms co produced to produce chaotic excitations. It is known that two coupled oscillators, a double pendulum, can produce chaotic behaviour.

The second way is if the surface network has defects. By defects we mean points or regions of surface discontinuity. If, for example, the axon is narrowed by scars or there are local periodic structure discontinuities due to connectivity changes then additional excitations can result. Such effects can be modelled by perturbations of the non-linear Schroedinger equation by impulses of short duration at these special locations or by introducing local periodic structure differences as potentials. It is known that chaotic excitations can be produced in the non-linear Schroedinger equation by perturbations of this type[68]‥

## 8 Conclusions

In this paper a surface network with special properties was studied. The network suggests testable solutions to a number of long standing issues in neuroscience research. The first issue is that although it is established that memories are stored in engrams, the questions: how are memories stored and in what form are they stored are unanswered. The surface model addresses this issue by suggesting that memories are aligned proton spins structures located in the myelin sheaths of axons. The axon pathways are between assembly of neurons and the memory structure they support is stable only if the pathways include loops.

The second issue is how are memories recalled. In the surface model the memory structure is a quantum spin system with each memory having a specific excitation frequency. Thus a given memory may be recalled by a resonance mechanism which excites it memory from an assembly of memories. This process was discussed and it was pointed out that it was similar to the way a quantum computer works.

The next issue is conceptual it suggest how an a system create its unknown communication code. The surface network suggests a way. The parameters of input pinch deformation signals that define the information content of a signal were shown to a self generated communication code.

The next issue is to explain the origin and predict the properties of EEG waveforms, namely why do they have five frequency bands. This issue was addressed by relating EEG waveforms to special oscillations on the surface of an assembly of neurons, induced whenever other brain signals are generated. The suggested mechanism correctly predict the observed frequency bands of EEG waveforms and remarkably relates them to the existence of the five Platonic solids of antiquity.

We now summarise some of the predictions of the approach. The most significant prediction is that memories are aligned proton spins, located in myelin sheaths of axons that belong to a cluster of neurons with non-zero spin topology number.

1. A strong prediction made was that soliton signals carry with them information regarding the way they are created. This information is encoded in the networks own code which is given by the unknown pinch deformation parameter values that generated the signal.
2. The example of thalamic neuron spikes discussed provides evidence that multi-soliton solutions of the non-linear Schroedinger equation can reproduce observed brain action potentials and the fits confirm the theoretical prediction that each spike is different.
3. We expect that action potentials from the different sensory organs that look the same are different in detail and expect that their nature (visual or auditory) can be determined from these details because they are created by different types of pinch deformations. Using machine leaning methods it should be possible to test this prediction.
4. The oscillation frequency bands and amplitude values of EEG waveforms as special sphere surface oscillations were predicted and were found to be in reasonable agreement with observations. Recent observations support the idea of neuron soma oscillations when action potentials are emitted. It is observed that the neuron soma is deformed when the neuron emits an action potential. The sequence of deformations observed was successfully modelled by treating the neuron soma as a sphere [34]. The observed surface also has ridges that may represent localised tiling waveforms.
5. In a recent paper by Pang et al[69], were trying to find the best global marker of brain activity. They choose solutions of the equation,∇2*ψ*_*λ*_ =−*λ*^2^*ψ*_*λ*_, where the Laplacian operator ∇2 is that of a sphere surface and found that these solutions provided the best marker for brain activity. This result confirms the expectation of the network. In the network EEG waveforms are special solutions of ∇2*ψ*_*λ*_ = −*λψ*_*λ*_ and they are predicted to be the best markers of brain activity as they are co-produced whenever any brain signal is created by topological means.
6. Since EEG oscillations are suggested to be due to oscillating neuron soma surfaces, they should be the same for all creatures with neurons. This prediction is confirmed for a very wide range of vertebrates, including fish, amphibians, reptiles, birds and mammals, large and small by Bullock[72] and for human beings with different mental disorders by Fingelkurts[73].
7. It is predicted that EEG waveforms are modulated by other brain events and these modulations themselves can lead to new excitations. We used these ideas to interpret the sequence of EEG modulations that follow a K-complex event, as a process of memory consolidation and transfer[5]. The special feature of this analysis is that it includes interactions between memories and incoming signals via EEG waveforms in a feedback loop.
8. The presence of spin structure is an essential part of the network dynamics. Pre-existing spin structures of the brain are expected to be present from birth. Hence memory formation, that require modifications of spin structure, do not start from a blank slate, but builds on an pre-existing structure[74].
9. The memory structure is predicted to have a natural, theoretically estimated, frequency and it is also predicted that a second associated memory structure with double this excitation frequency should be present due to memory structure spin pairing. Such a double frequency has been experimentally observed[75]. This frequency doubling is important as it means low frequency gamma waves can also excite the memory[33].

The method of EEG waveform generation proposed here differs from those currently proposed[2, 76, 77, 78, 79, 80] in an essential way. Here EEG waveform creation is directly related to a coherent process of surface oscillations of a large number of neurons, each producing a mix of the distinct classes of surface oscillations discussed, whenever excitations are generated in response to pinch deformations described in this paper. But in all existing theories action potential generation and EEG waveforms are not directly related.

### 8.1 Future Directions

The focus of the paper was on a method of generating all brain-like signals. As we proceeded it became clear that the approach naturally suggested a way for storing memory and it suggested that the brain may operate using a small number of elementary processes, each one of which is shown in the network to have a mathematical representation. Underlying all of them is the one key process: the selective blocking of signal pathways. In neural field theory this fundamental feature is built in as the balance between inhibitory and excitatory neurons for each brain process. In the network we need to find a global mechanisms that does this. In our discussions we introduced transient localised excitations as a way of doing this. Such signals exist and can be generated by pinch deformations and they are thus globally generated but what triggers them needs to be understood. We suspect the answer lies in the trigger signals induced by hormone/chemical/gas releases in the brain. These blocking events allow focused attention and are thus vital for all cognitive functions of the brain.

In view of these remarks the simulations of brain activity in the network should involve a sequence of linked elementary brain processes each of which has a mathematical representation. The path of activity is expected to be driven by the environment and the intentions of a person and will involve thresholds set by feedback loops of interactions between memory and signals and between signals and hormone/chemical releases. These feedback loops are space-time dependent. A sketch of how such an approach would work was given in the description of the K-complex sequence of events. But a general operational scheme is lacking.

The immediate challenge is to observationally confirm or rule out our suggested structure for memory and its conjectured location. The current work is the start of a new global way of thinking about the brain that might be useful.

## Author Contributions

The author carried out all the formal analysis and wrote the paper. The work is theoretical.

## There is no new data used in the paper

The paper is theoretical and makes use of published data. No new data is used in the paper.

## Ethics

No ethical issues arise as the work is theoretical.

## Use of AI

No AI tools were used.

## Methodology and Resources

The methodology used is built on earlier theoretical work done in collaboration with Tomas Ryan, Maurizio Pezzoli, Plamen Stemanov and David Muldowney that is referenced and is publicly available. Resources used are published papers and those in the public domain.

## Acknowledgement

I would like to thank Tomas Ryan and Maurizio Pezzoli for many discussions, David Muldowney for collaborating in the early stages of this work, Samik Sen for producing all the EEG, K-complex and sleep spindle plots using GnuPlot software, Maurizio Pezzoli for producing some figures using public domain software and Mani Ramaswami for encouragement. The data and fits of thalamic neuron spikes reproduced are due Maurizio Pezzoli of EPL, Switzerland (data) and Planem Stemanov of Trinity College Dublin (fits). Finally I would like to thank Mike Coey, for producing the figure of the surface connectome, for carefully editing the text and suggesting that proton spins might underlie memory structures.

## Supplementary Information

### Previato’s dark N-Soliton Solution

The dark N-soliton solution found by Previato[27]was used to fit the data of Pezzoli by Stemanov.

### Previato’s Dark Soliton Solution Details

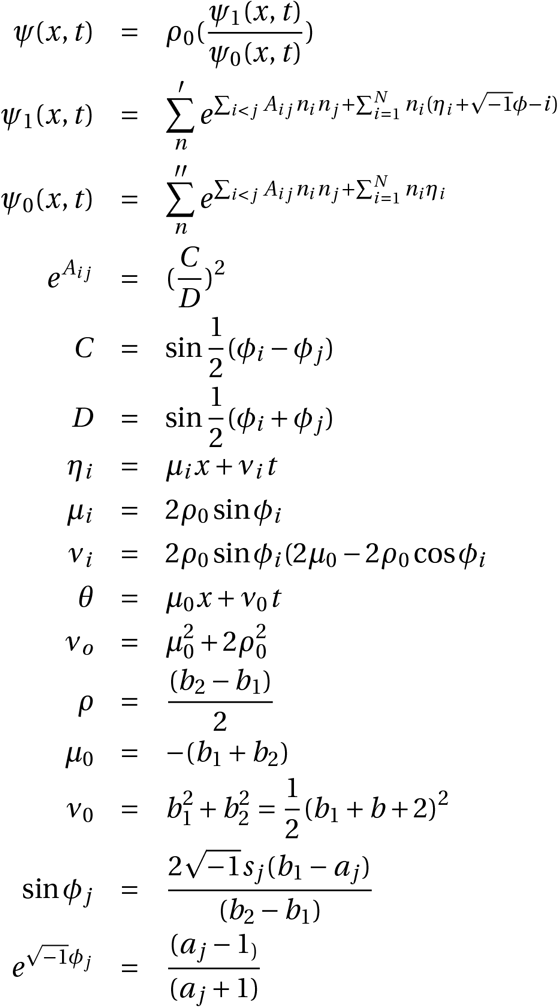

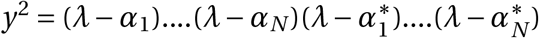 equation defining Riemann surface.

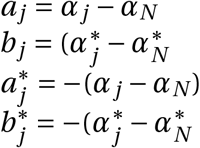

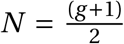 We assume *g* is odd. *N* is the number of spikes and 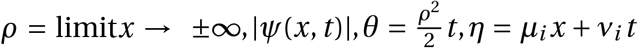. The parameter values for *A, µ, ν, χ* depend on pinch deformation parameters. The sum over 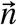 is for all values of the *n*_*i*_ that are either *n*_*i*_ = 0, 1 with 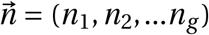 and *N* = *g*. Now the boundary condition is explicit and there is no oscillations in the *x* variable. The single time oscillation terms is an overall multiplicative factor.

### Spin Memory Structure has no Macroscopic Magnetism

We will describe how the memory structure is formed and show that as a result of the special topological feature of spin half particles under 2*π* rotation the structure created has no net macroscopic magnetic properties.

The non transient helical spin structure is determined by the quantum time evolution operator 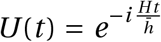, where 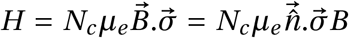 where we have written 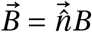 and 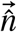 is the direction of the magnetic field 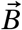 and *µ*_*e*_ = 2*µ*, where *µ* is the magetic moment of a proton. The magnetic field interacts with *N*_*c*_ spins at each point *z* that are on a circle round the axon tube with centre *z*. We now use the identity, which follows from the fact that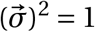.

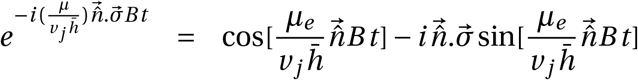

The time evolution of an initial spin *S* _*j*_(0) located at a point, is given by the equation,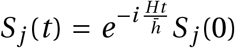. In our case their are *N*_*c*_ proton spins on a circle round the axon tube at any point *z* which is the location of the transient magnetic field pulses. These pulses aligns the spins. To study the nature of the cluster of *N*_*c*_ aligned spin distribution, parametrized by z, we replace *t* by 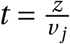, where *z* is a surface point on the pathway of the transient magnetic field and *v* _*j*_ is the speed of the pulse. Then we have the spatial spin aligning evolution operator *U* (*z*), given by,

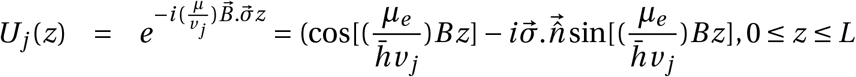

We write 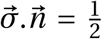, which is the aligned spin value projected out by the evolution operator, and replace *µ*_*e*_ by 2*µ* the Bohr magneton. Our static helical spin-magnet distribution,*S* _*j*_(*z*) is given by,

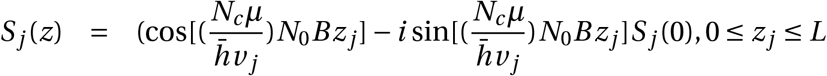

It is a helical chain of surface electron spins, where *S* _*j*_(0) represents the aligned spin at *z* = 0. It is created by the transient magnetic field pulses of a single soliton pulse. For it to store memory it has to be protected against body temperature thermal disordering by electron spin interactions[2]. There are three numbers that decide if a memory can be formed and if it is stable. The first number *N*_*j*_ is the charge carried by the *j* ^*th*^ soliton pulse, the second is the number of spins present on a circle on the axon tube, *N*_*c*_ and the third number id *N*_*s*_ the number of soliton pulses in the memory creating spike train.

We next note that the spin memory structure does not produce a macroscopic magnetization as two spin magnets at neighbouring sites on the helix, related by 2*π* rotation, have opposite magnetic directions since a spin changes its direction after a rotation of 2*π*. To analyse spin dynamics quantum methods have to be used[29, 30]. As a result the memory structure is helical, is based on spin magnetism but it does not have a macroscopic magnetic field as neighbouring spins are oppositely aligned. For *N*_*s*_ pulses there will be *g* static structures given by 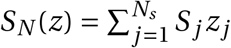. For estimates we set *N*_*j*_ = *N*_0_, *B*_*j*_ = *B, S* _*j*_(0) = *S*(0) We write this as,

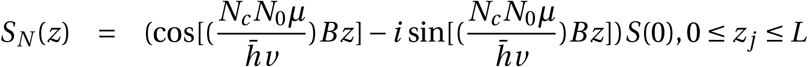

We next show that since spins at adjacent helical sites with related by 2*π* rotations have opposite orientations, they pairing form zero spin, charge 2*e* objects with no net magnetism. These charged electron pairs are stable in an environment with ions present. Spin zero objects are bosons and are not prevented from coming closer together by the Pauli principle. Consequently the oppositely oriented spins can come closer together thus increasing their binding. This paired spin structure defines a second effective memory copy. We write the structure in terms of the correlated pair of electron spins function, < *C* (*z*) >=< *S*(*z*)_↑_*S*(*z* +*a*)_↓_ > where the pair represent neighbouring helical sites, *z, z* + *a*. For small *a* a simple expression for the helical memory structure in terms of < *C* (*z*) > can be obtained, given by,

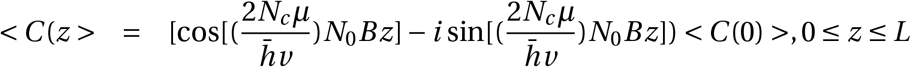

### Estimates of EEG frequencies and Amplitudes

We next give a simple estimate of the Frequency/Amplitude scales of for EEG waveforms by relating the energy of oscillation of a point on the surface of a sphere to the maximum voltage oscillation energy of EEG waveforms observed. This fixes the frequency scale of the system and relates it to the amplitude scale. But to get a precise expression for allowed frequencies and the corresponding EEG waveform shapes we have to use the tiling solutions of the wave equation on the surface of a sphere. Now we have a precise formula for allowed frequencies but the scale of frequencies is not fixed by the equation. We use our simple estimate to choose an appropriate realistic scale and complete our analysis.

The energy of an electron located at a point on a spherical surface, oscillating transverse to the neuron surface, with velocity *v*, can be written as, 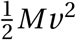. We set this equal to the voltage energy *eV*. Since the oscillations are coherent we set *M* = 4*πρR*^2^, the surface mass, with *ρ* the surface density. We then write the square of the oscillating points speed as *v* ^2^ = *ω*^2^Δ^2^ where *ω* is its frequency, *ρ* the average density of the surface and Δ the maximum amplitude of the surface oscillations. Setting 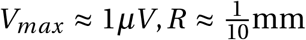 and *ρ* ≈ 1 gm/cc (estimate), we predict Δ < 10^−6^cm and confirm that the frequency is inversely related to the frequency.

The oscillation frequencies of the neuron surfaces determined have voltage amplitudes greater than those observed on the scalp. In our discussions we will assume a linear scaling rule for converting neuron surface oscillations amplitudes that are in milli volts[1], to the micro volt amplitudes observed on the brain scalp.

We can also estimate the speed range of the different EEG waveforms *v*, since *v* = *λω* ≈ *R* × *ω* ≈< 1cm/s. There will be range of speeds close to this estimate. The higher frequency waves have shorter wavelengths but higher frequencies they should also have speeds in the same range. These estimate show that frequencies allowed will be in bands as observed EEG waveforms are due to signals from a large number of spheres created by a large number of soliton pulses and different spheres created will have different radii and densities.

We can also estimate the speed of soliton pulse signals inside the soma by using a conservation of angular momentum argument. Suppose the pulse is moving in the *z* direction with speed *v*_*s*_ in the soma that has a radius *r*_*s*_. When it enters the pinched axon hillock its speed is taken to be *v*_*a*_ and the pinched hillock radius is taken to be *r*_*a*_. Angular momentum conservation requires *v*_*s*_*r*_*s*_ = *v*_*a*_*r*_*a*_. Setting *v*_*a*_ ≈ 10^4^*cm*/*s, r*_*a*_ ≈ 10^−5^*cm, r*_*s*_ ≈ 10^−2^*cm* we get *v*_*s*_ ≈ 10*cm*/*s*.

### Construction Greens function

In this Appendix we construct the Greens function appropriate for dihedral tiling solutions of the wave equation on the surface of a sphere. The wave equation Greens function *G*(*z, t* : *z*^′^, *t* ^′^) satisfies the equation,

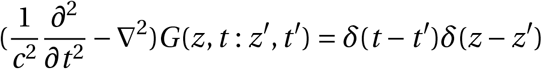

Once the Greens function is known it can be used to find the distribution of dihedral tiling waveforms generated by any input signal. Our starting point is the result of Maier[45]who showed that the associated Legendre functions of fractional order, 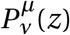, form a complete set of basis functions. This means that they can be used to represent any arbitrary square integrable function *f* (*z*) on the sphere. Thus, any such function *f* (*z*), on the surface of the sphere can be written in terms of tiling solutions as,

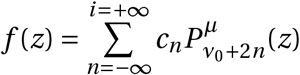

where the points *z* represent arcs on the sphere. From this result and the orthogonality property,

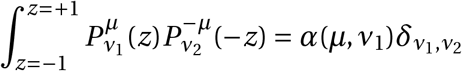

we obtain the completeness result,

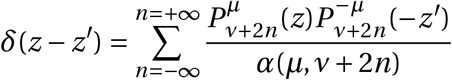

The dihedral tiling class corresponds to setting 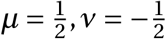. We also have the completeness result,

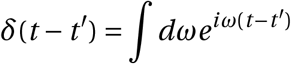

We now use the completeness property of these dihedral tiling solutions of the wave equation to construct a dihedral tiling Greens function *G*(*z, t* : *z*^′^, *t* ^′^) [3]by replacing the delta function present in the Greens function equation by its representations in terms of dihedral tiling solutions, where we set 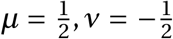. Thus we write,

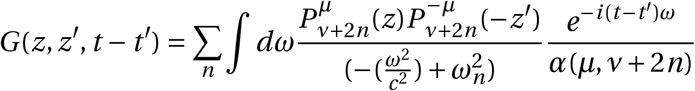

where 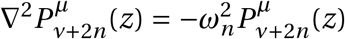 and we have assumed that the Greens function depends only on (*t* − *t*). Carrying out the *ω* integral we get our Greens function,

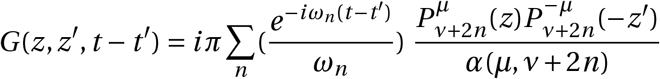

### Two practical uses of EEG waveforms

We next discuss two practical consequences of the model that illustrate the importance of EEG waveforms. The first uses the mathematical result that EEG waveforms, corresponding to dihedral tiling form a complete basis so that they can be used to represent any arbitrary function, representing an electrical event of the brain. The dihedral waveforms, written down earlier Eq(1), are given by,

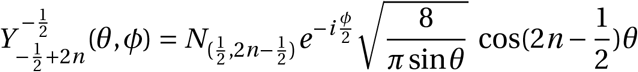

where *n* = 1, 2, ‥ and 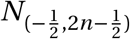 is a normalization factor. Thus any observed brain excitation distribution *O*(*θ*.*ϕ*) can be written as a superposition of the dihedral waveforms[45]. We can write,

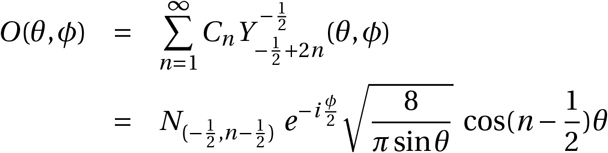

where *n* = 1, 2, ‥ and 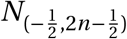 is a normalization factor, given by 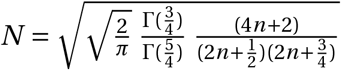 The coefficients can be determined as integrals of *O*(*θ, ϕ*) multiplied by the complex conjugate of the basis tiling waveforms. Namely,

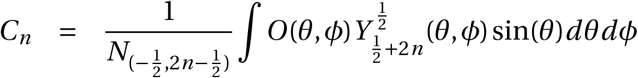

Using these two expressions any observed brain excitation can be decomposed into modulated EEG waveforms. Since the suggests that EEG waveform are co produced whenever any brain excitation is generated, we would expect that EEG waveforms,*ψ*(*x*)_*λ*_ eigenstates of the sphere Laplacian equation, ∇2*ψ*(*x*)_*λ*_ = −*λ*^2^*ψ*(*x*)_*λ*_, where *x* is a sphere coordinate, they will be a good markers of brain activity. This prediction has been confirmed[69].

The second application of EEG waveforms addresses the reverse question: Given an observed brain surface waveform *R*(*θ, ϕ, t*) can one find the source signals *S*(*θ*^′^, *ϕ*^′^) that generated it?

This can be done by expressing *R*(*θ, ϕ, t*) in terms of the complete set of time dependent dihedral tiling EEG waveforms. Each tiling waveform appearing can be related to the size of active brain circuits involved in producing the source signal since we showed earlier that both the size of the subunit and the nature of a EEG waveform are fixed by the integer value of the spin topology number *W*. Thus each EEG waveform gives information about the *W* that generated it and the *W* value is a measure of the circuit size involved. The modulations of the tiling will provide additional confirmation of the nature and location of the source event.

Consider an example. Suppose we find the presence of delta waveforms in the source. Our theoretical result then tells us that a delta waveforms, of frequency 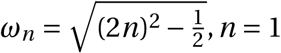 is generated by three input solitons with spin topology numbers *t*_*i*_ given by 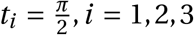 This excitation corresponds to the excitation of a localised brain circuits of low genus (g=2), while for gamma waveforms of frequency *ω*_*n*_, *n* > 20 the phases are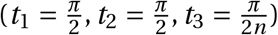, thus larger brain circuits are involved. The procedure described should help locate the brain circuits and the neurons responsible for an epileptic episode.

## Footnotes

* We summarise some of the results of our paper[2]that we need to discuss EEG waveforms and the way they interact with memory

† The brain connectivity architecture evolves. Thus a connectome changes. Our result hold for a changing connectome as well

‡ When neurons produce action potentials it is observed that their surface is deformed to a more spherical shape and they have surface deformations[34].

§ The magnetic field generated by an action potential has been measured and has been used to accurately reconstructed the original voltage pulse signal[25]. Thus encodes all the information carried by the signal.

¶ Helical spin half structures suggested here are possible, and have been observed in condensed matter physics[26].

|| The technical details of the deformed Fay identity are omitted. They are discussed in a separate paper[2].

** The argument is due to Mike Coey, private communication

†† The pairing of neighbouring spins makes the pair a boson so that they can cluster together unaffected by the Pauli exclusion principle repulsive effects. Thus increases the stability of the structure.

‡‡ There is experimental evidence that the soma of neurons become more spherical when they emit action potential and have observed surface ridges that have been measured[34].

